# Soft-Robotic Magnetic Microfluidic Catheter for Delivery of Aqueous-Based Dual-Component Embolic Formulations

**DOI:** 10.64898/2026.09.14.751422

**Authors:** Michael Mattmann, Carlos Franco, Hao Ye, Fabian Landers, Lucas Hertle, Peter Fischer, Nicole Ochsenbein, Maja Ruetten, João Pedro Vale, Jonas Lussi, Josep Puigmarti, Tiago Sotto Mayor, Pedro D. Wendel-Garcia, Silvana Geleff, Kamal Farha, Said Farha, Philipp Gruber, Miriam Weisskopf, Salvador Pané, Bradley Nelson

## Abstract

Transcatheter embolization requires materials that can be steered through tortuous vessels, solidify rapidly in situ, remain clearly visible under fluoroscopy, and, ideally, carry therapeutic cargo without harming tissue. To meet these requirements, we present a fully water-based, two-component PEI-PEG hydrogel delivered through a soft-robotic, microfluidic catheter that keeps the precursors separate until they meet in a millimetre-scale mixing chamber at the tip. Fast amide cross-linking converts the liquid pair into a self-supporting gel within seconds, eliminating organic solvents and preventing catheter blockage. By adjusting precursor ratio and flow regime, the gel’s stiffness and viscosity can be tuned over orders of magnitude, with the same chemistry allowing to occlude both high-flow arteries and fragile micro-vessels. The platform was validated in three escalating models. First, in ex-vivo perfused human placenta, the hydrogel filled targeted branches without reflux or fragmentation, demonstrating controlled delivery in clinically relevant vasculature. Next, in three porcine embolizations, splenic, hepatic and ascending pharyngeal arteries, the material achieved stable, selective occlusion with no migration, vasospasm or recanalization, showing seamless compatibility with standard interventional workflows. Finally, in rats bearing orthotopic liver tumours, drug-loaded hydrogel delivered through the hepatic artery concentrated doxorubicin inside tumours while sparing healthy tissue, confirming its potential for precision chemoembolization. These results position the PEI-PEG hydrogel and microfluidic catheter as a unified, image-guided platform that couples robust mechanical occlusion with site-specific drug delivery, offering a biocompatible alternative to current liquid embolics and expanding the therapeutic reach of minimally invasive embolization procedures.

## 1 Introduction

Transcatheter embolization (TCE) is a minimally invasive procedure that involves the deployment of (bio)materials through a catheter into a specific region within a blood vessel^1^. The primary aim of embolization is to block the blood flow to a specific area of the body for treating an aneurysm, eliminate abnormal vessel connectivity or malformations, or starve an unresectable tumour from its blood supply^2–5^. Embolic materials can also serve as platforms for the localized delivery of therapeutic agents, providing a targeted approach that improves treatment efficacy^6,7^. Compared to conventional surgery and systemic drug administration, TCE offers several advantages, including minimally invasive access, reduced systemic toxicity, localized therapeutic delivery, and lower overall treatment costs^5,8^.

Current embolic agents, including coils, foams, flow disruptors, and microparticles, suffer from key limitations such as particle migration, fragmentation, poor biodegradability, and mechanical instability, as well as reduced shelf life, low traceability, and restricted drug-loading capacity^9,10^. These shortcomings hinder their adaptability across clinical scenarios and limit their use in advanced therapies. Liquid embolic agents, such as cyanoacrylates and ethylene-vinyl alcohol copolymers, have emerged as promising alternatives due to their ability to conformally fill vascular structures and deliver therapeutic payloads independently of a patient’s coagulation status^11,12^. However, their broader adoption is hampered by critical challenges in biocompatibility, injectability, and imaging^13,14^. Many formulations require organic solvents to reduce viscosity, which compromises safety and prevents integration with emerging biotherapeutic cargoes such as proteins, enzymes, or cells^15,16^, while other formulations transition to gel-like states that may obstruct catheter tips or complicate repeated injections^17^. Most of these lack intrinsic radiopacity, hindering real-time imaging of the emboli placement and persistence^18^. Moreover, catheter compatibility remains a major constraint: materials that tolerate harsh solvents require stiffer catheters, reducing their maneuverability in deep, tortuous vessels^9^. Finally, navigation itself presents a formidable challenge. Increased vascular friction and poor trackability can lead to catheter misplacement or vessel trauma, while attempts to improve steerability often sacrifice flexibility, intensifying these risks^19^. Collectively, these issues underscore the pressing need for a next-generation embolic platform that can couple precise, image-guided delivery with versatile drug loading, high stability, and seamless catheter integration^9,19^.

To address these challenges, we introduce a biocompatible, in situ forming PEI-PEG hydrogel system tailored for embolization, delivered through a microfluidic-assisted catheter platform and validated in three biologically relevant models. The hydrogel is based on the spontaneous amide crosslinking of succinimidyl-activated PEG (SPA-PEG-SPA) with dendritic polyethyleneimine (PEI), producing a highly viscous, self-supporting network within seconds of mixing. Our design enables precise control over gelation by separating the reactive components until mixing at a microfluidic coaxial chamber at the tip of the catheter. This setup ensures clog-free delivery, fast setting, and customizable mechanical properties. We benchmarked our approach in three models of increasing complexity: (i) ex vivo perfused human placenta vasculature, which allows direct visualization and quantification of embolic performance in a clinically relevant scale; (ii) in vivo embolization of uterine arteries in a porcine model, simulating a real surgical workflow; and (iii) intra-arterial catheter-based embolization in rats bearing orthotopic liver tumors, used to demonstrate targeted intra-tumoral drug delivery. In the placenta, we achieved stable occlusion of targeted vascular branches using <0.3 mL of hydrogel with no evidence of reflux. In pigs, selective embolization was successful in 3/3 cases, with complete vessel occlusion maintained over the surgical window. In the rat model, the hydrogel was loaded with a chemotherapeutic agent and delivered via catheter directly into tumor-feeding hepatic arteries. Post-treatment analysis showed localized drug accumulation in tumor tissue and minimal off-target distribution, confirming the hydrogel’s potential for precision chemoembolization. These results highlight the translational potential of our platform, offering a mechanically stable, non-toxic embolic material with programmable gelation kinetics and integrated drug delivery capabilities for catheter-based oncologic interventions.

**Fig. 1.**
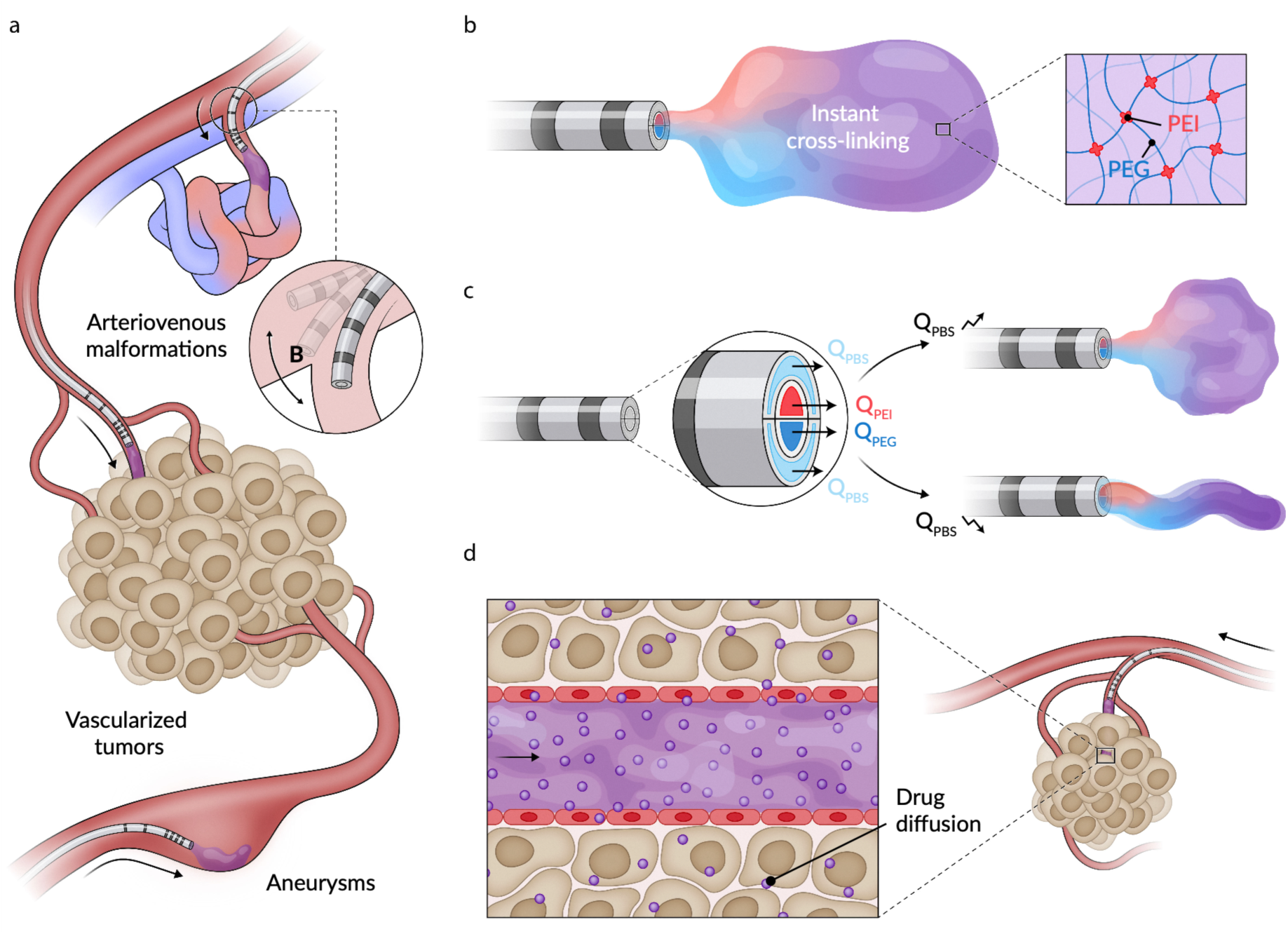
Application and design of embolic material and delivery system. **a**, Possible applications of the embolic system. The magnetic catheter system provides improved navigability with the ability to locally inject a tunable embolic material. **b**, The embolic material is injected through a microfluidic catheter and crosslinks immediately when mixed. **c**, The microfluidic catheter design allows for the controlled tuning of the embolus mechanical and morphological properties. **d**, Drug and protein loading capabilities allow for the localized drug delivery with possible applications in chemoembolization.

## 2 Results

### 2.1 Embolic material

Polyethylene glycols (PEGs) are hydrophilic and can form a wide range of compounds and complexes. These compounds are known for their water solubility, non-toxicity, and non-immunogenicity, making them highly suitable for various biomedical applications^20,21^. Notably, the FDA has approved several PEGylated drugs for the treatment of chronic diseases^22,23^. Moreover, PEGs demonstrate exceptional versatility as reagents, facilitating bioconjugation, crosslinking, and interactions among macromolecules, therapeutic compounds, and dyes^24,25^. PEGs have also been extensively studied as hydrogels for applications such as wound healing, tissue engineering, and drug delivery due to their tunable physical properties and biocompatibility^26,27^. Building on these characteristics, here we focus on exploring the potential of PEG derivatives as water-based vascular embolization hydrogels, with a specific focus on activated PEG. Activated PEG is a PEG whose chains were modified with reactive activated esters capable of reacting with nucleophilic crosslinkers^28,29^. For our research, we have selected succinimidyl propionate-polyethylene glycol-succinimidyl propionate (SPA-PEG-SPA) as our activated PEG derivative, specifically utilizing SPA-PEG-SPA of medium molecular weight of 3,400 g/mol. This compound features two non-toxic and biocompatible activated succinimidyl esters located at the ends of linear PEG chains^30,31^. To create the PEG hydrogel, we have selected polyethyleneimine (PEI) as the crosslinker. PEI, a commonly used biomedical agent, contains primary amines that readily react with the activated succinimidyl esters to form the PEG-based hydrogel^32^. Specifically, we opted for dendrimeric PEI with a low molecular weight (MW: 1200 g/mol) due to its reduced cytotoxicity compared to higher molecular weight counterparts and an appropriate number of reactive primary amines (Scheme 1)^33^.

**Scheme 1:**
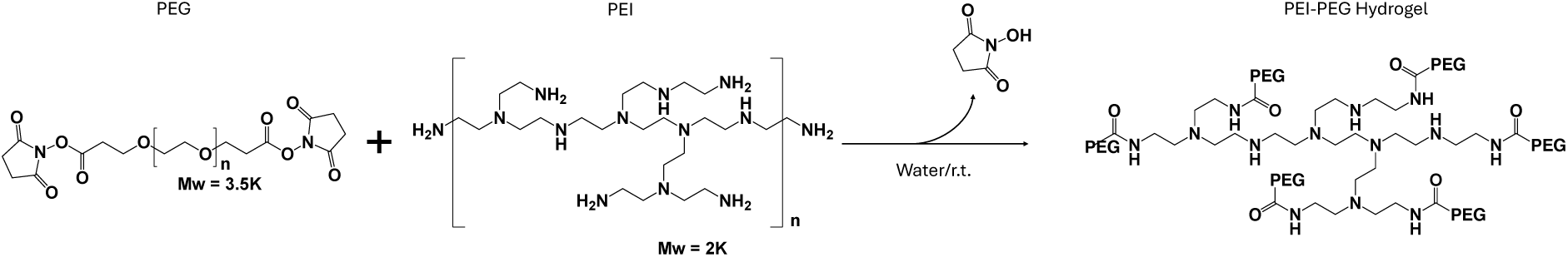
Schematic synthesis of PEI-PEG hydrogel

In an initial bench-top test, we dispensed equal volumes of SPA-PEG-SPA (332 mg ml⁻¹, PBS buffer) and dendritic PEI (20 mg ml⁻¹ PBS buffer) side-by-side in a 35 mm Petri dish and gently drew the two droplets together with a pipette tip. As soon as the fronts met, a translucent filament formed and spread laterally, converting the mixture into a self-supporting gel within ≈ 3 s (**Supplementary Video 1**). Tilting the dish immediately afterwards showed that the material no longer flowed, demonstrating a change from a low-viscosity solution to a highly viscous network. These qualitative observations confirm that primary amines on PEI rapidly crosslink the activated PEG chains under physiological conditions and lay the groundwork for the quantitative mechanical studies.

### 2.2 Rheological and mechanical properties

Because the rheological properties of the hydrogel network depend on the interaction between PEG and PEI, we have adopted a microfluidic approach to precisely control the reaction between the two components of the embolic formulation. In microfluidic environments, mass transport occurs mostly by diffusion, which allows for exceptional control over the mixing and reaction conditions and is essential for achieving consistent properties in the resulting PEI-PEG hydrogel^34^. To leverage the enhanced control offered by microfluidics for the controlled formation and delivery of the PEI-PEG hydrogel, we designed a sub-millimeter microfluidic-like catheter (outer diameter 0.9 mm) made in polyurethane (see specifications in **Supplementary Fig. 1b**), with a design featuring a 3D focusing microfluidic approach based on four inlet lumens (see **Supplementary Fig. 1a,b**). The two central lumens deliver PEG and PEI solutions, while the two external lumens deliver the solvent (PBS) that is used to regulate the flow focusing. These streams converge in a single 1-cm long outer lumen that acts as a microfluidic reactor, providing a controlled reaction-diffusion (RD) environment for the precise mixing of PEG and PEI solutions. We performed 3D numerical simulations of flow and mass transport in our microfluidic catheter device to investigate the mixing of PEI and PEG. We confirmed that the reactants mix primarily by diffusion, that a RD zone is formed between the reacting streams where most of the embolic hydrogel is generated, and that the RD zone grows in width along the microfluidic device (**Supplementary Fig. 10**). The laminar flow (Re ≈ 20) in our microfluidic setup enables us to maintain this RD zone unchanged throughout an experiment and, as will be demonstrated, facilitates the fine-tuning of the hydrogel’s mechanical and morphological properties by simply modulating the flow rates.

To optimize the design of the microfluidic device, we compared the RD zone generated in different flow-focusing setups (**Supplementary Fig. 11**). Specifically, we compared the RD zone generated using the proposed multi-lumen setup (**Supplementary Fig. 11a**) with that generated using two commonly used microfluidic flow-focusing setups: a 2D planar design (**Supplementary Fig. 11b**) and a co-axial design (**Supplementary Fig. 11c**). If we were to employ the 2D planar design (**Supplementary Fig. 11b**), the RD zone would be in contact with the walls of the device, leading to uncontrolled delivery of the hydrogel and clogging caused by attachment of the fibrous structure to the walls. Alternatively, if we were to employ the co-axial design (**Supplementary Fig. 11c**), the RD zone would form a ring-shape which would lead to the generation of a hollow fibrous structure that would not be ideal for the complete occlusion of vascular structures. In contrast, using the multi-lumen design proposed here (**Supplementary Fig. 11a**), the RD zone does not contact the walls of the device and is not hollow, allowing for greater control over the delivery of the hydrogel and for optimal embolization performance.

In a typical experiment, we injected PEG (332 mg/mL) and PEI (20 mg/mL) with an equal flow rate of 300 µL/min to obtain the PEI-PEG hydrogel. Rheological studies performed on the PEI-PEG hydrogels that were generated under these microfluidic conditions indicate that the materials exhibit a solid-like elastic behaviour, with the storage modulus (G’) consistently exceeding the loss modulus (G’’), as shown in **Fig. 2a**. Additionally, the viscosity of the hydrogels decreased with increasing oscillation frequency (**Fig. 2b**), indicating shear-thinning behaviour. This type of rheology is ideal for an embolic material, as the hydrogel will have a low viscosity (liquid-like) during delivery and a high viscosity (solid-like) when occluding vessels. To assess the recoverability of the hydrogel under cyclic high and low shear strain, thixotropy tests were conducted. These studies revealed that the PEI-PEG hydrogels experienced a drop in storage modulus at high strain but recovered its original modulus when the shear strain decreased, demonstrating stable thixotropic behaviour (**Fig. 2c**).

**Fig. 2.**
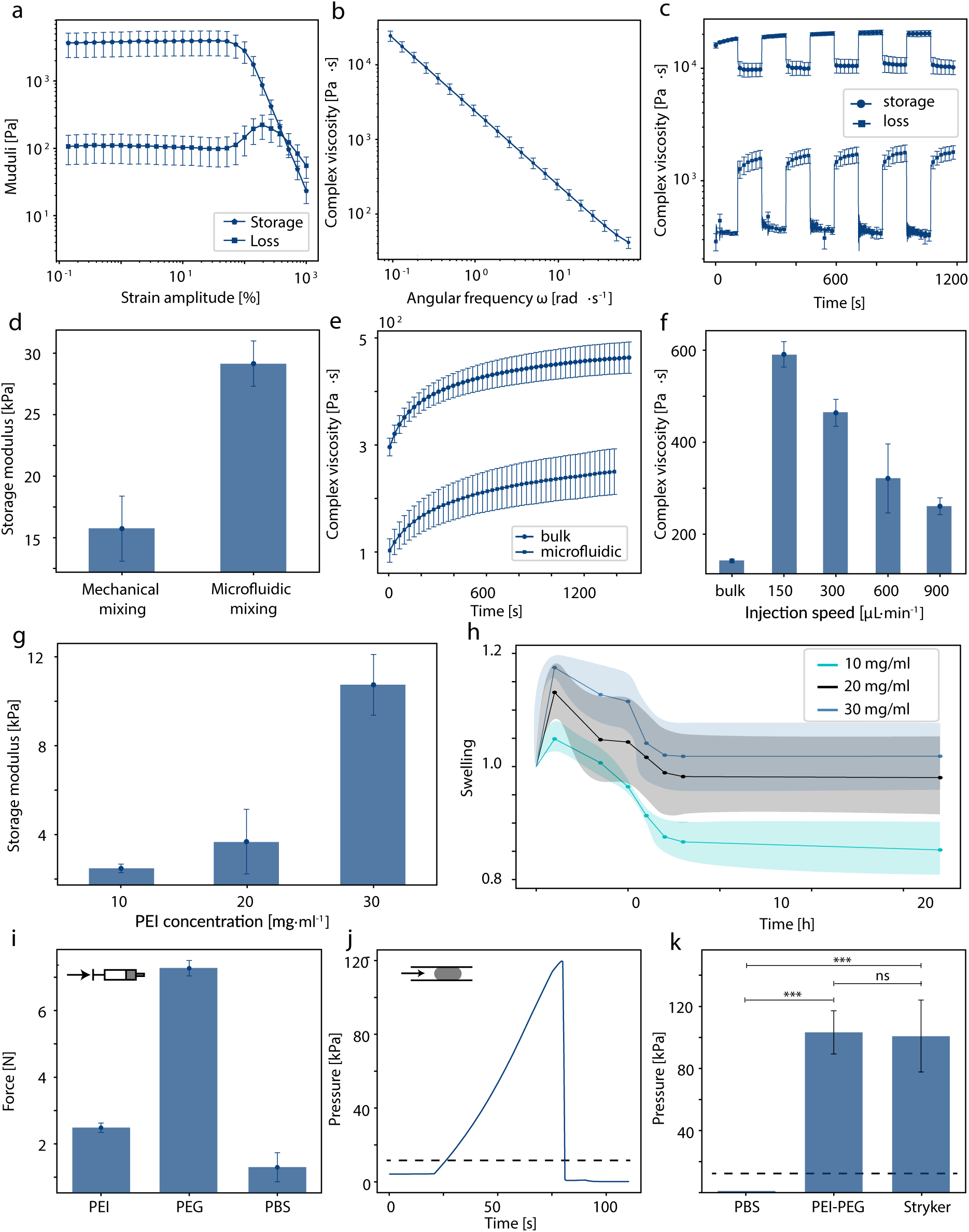
Mechanical characterization of PEI-PEG hydrogels containing 40 wt% of Histodenz. **a**, Rheological strain amplitude sweep showing a solid-like behavior. **b**, Frequency sweep of a PEI-PEG hydrogel showing a shear thinning behavior. **c**, Thixotropic behavior of PEI-PEG hydrogels to assess recoverability and fragmentation under high-low load cycles. **d**, Storage modulus of hydrogels generated with mechanical and microfluidic mixing. **e**, Complex viscosity over time shows the crosslinking speed for microfluidically and mechanically mixed hydrogels. **f**, Complex viscosity with a varying injection speed of PEI and PEG. **g**,The storage modulus of PEI-PEG hydrogels with varying concentrations of PEI. **h**, Swelling properties of PEI-PEG hydrogels at different PEI concentrations. **i**, Injection force of PEI, PEG, and PBS passing through a 140 cm long microfluidic catheter at a flow rate of 300 µL/min, 300 µL/min, and 600 µL/min, respectively. **j**, Representative pressure curve to displace a PEI-PEG hydrogel. **k**, Pressure required to displace PBS (control), PEI-PEG hydrogel, and Adherus AutoSpray hydrogel. Statistical significance was determined using one-way ANOVA. ns, not significant; \**p* < 0.05, \*\**p* < 0.01, and \*\*\**p* < 0.001.

To investigate the influence of the mixing strategy on the material properties, we compared the rheological properties of hydrogels generated under microfluidic conditions versus those obtained by mixing in bulk. The materials injected using microfluidics showed a two-fold increase in complex viscosity (**Supplementary Fig. 2d**) and storage modulus (**Fig. 2d,e**) compared to the values observed in the bulk-mixed samples. This suggests that greater entanglement between polymer chains is achieved at the microstructural level, leading to higher stiffness in the resulting hydrogel. Furthermore, the injection flow rate significantly influenced the complex viscosity of the hydrogels generated using microfluidics (**Fig. 2f**). Higher flow rates originated hydrogels that had lower complex viscosity, approaching the values of bulk-mixed hydrogels, while lower injection speeds promoted increased crosslinking within the mixing chamber, leading to an increase in the complex viscosity of the injected material.

The relative injection rates (i.e., the central-to-sheath flow-rate ratio) also influenced the morphology of the PEI-PEG hydrogel. Identical rates produced a symmetric flow field with uniform mass transport, yielding stable and continuous filaments. In contrast, mismatched rates introduced shear asymmetries that disrupted the flow and led to irregular, wall-anchored structures. This behaviour was examined through numerical simulations performed under different flow conditions (**Supplementary Fig. 12**) and verified experimentally (**Supplementary Fig. 1c**). The simulations revealed that, when flow focusing is lost, the RD zone expands until it contacts the channel wall (**Supplementary Fig. 12c**), which could initiate gelation at the boundary and lead to complete occlusion of the microfluidic device (**Supplementary Fig. 1c**). Conversely, when equal injection rates are used, the RD zone remains centred and isolated from the channel walls (**Supplementary Fig. 12a**), allowing for the controlled formation of continuous hydrogel filaments (**Supplementary Fig. 1c**). This highlights the importance of generating the hydrogels in controlled environments for generating effective linkage points. Moreover, tuning the individual and relative injection speeds enables the modulation of the RD zone where hydrogels are generated, allowing for precise control over the mechanical and morphological properties of the injected material.

To further understand the influence of polymer concentration on the mechanical properties of the hydrogels, we analyzed a series of PEI-PEG hydrogels with varying PEI concentrations. An increase in PEI concentration led to a corresponding increase in both the storage and loss moduli (**Fig. 2c and Supplementary Fig. 2c**). Similarly, the complex viscosity increased with higher crosslinker concentrations (**Supplementary Fig. 2b**), though it eventually reached a plateau, indicating a limited influence on injectability. However, the swelling properties of the hydrogels were not significantly affected by PEI concentration. Higher concentrations resulted in slightly increased swelling, with marginal swelling observed in the initial hours that stabilized after approximately 5 hours (**Fig. 2h**). This indicates that while the hydrogel’s initial swelling is responsive to PEI concentration, the long-term stability remains consistent, suggesting that the hydrogels can maintain their structural integrity over time in a physiological environment.

### 2.3 Visibility

Injection trackability is crucial for the success of various endovascular interventions, as it provides direct feedback on the location and volume of the injected material^35^. During these procedures, X-ray fluoroscopy is commonly used to visualize the embolic material in real-time^36^. Therefore, high X-ray contrast and good visibility against high-contrast anatomical structures, such as bones, is essential for embolic formulations^5^. To enhance the visibility of our hydrogels under X-ray fluoroscopy, we introduced standard iodine-based contrast agents. Specifically, we used Histodenz as a contrast agent in our PEI-PEG hydrogels, which is easily soluble in water, thus allowing its inclusion in our hydrogel formula without affecting the hydrogel formation and stability. Fluoroscopic imaging of PEI-PEG hydrogels with a Histodenz concentration between 0 and 40 wt% showed a concentration-dependent increase in radiodensity (**Fig. 3a**). Importantly, the visibility of the hydrogels improved as the concentration of Histodenz increased, demonstrating a clear, progressive enhancement in radiographic contrast. Additionally, a PEI-PEG hydrogel with a 40 wt% contrast agent exhibited excellent visibility within an intracranial pig’s vessel under fluoroscopic imaging (**Fig. 6c**).

**Fig. 3.**
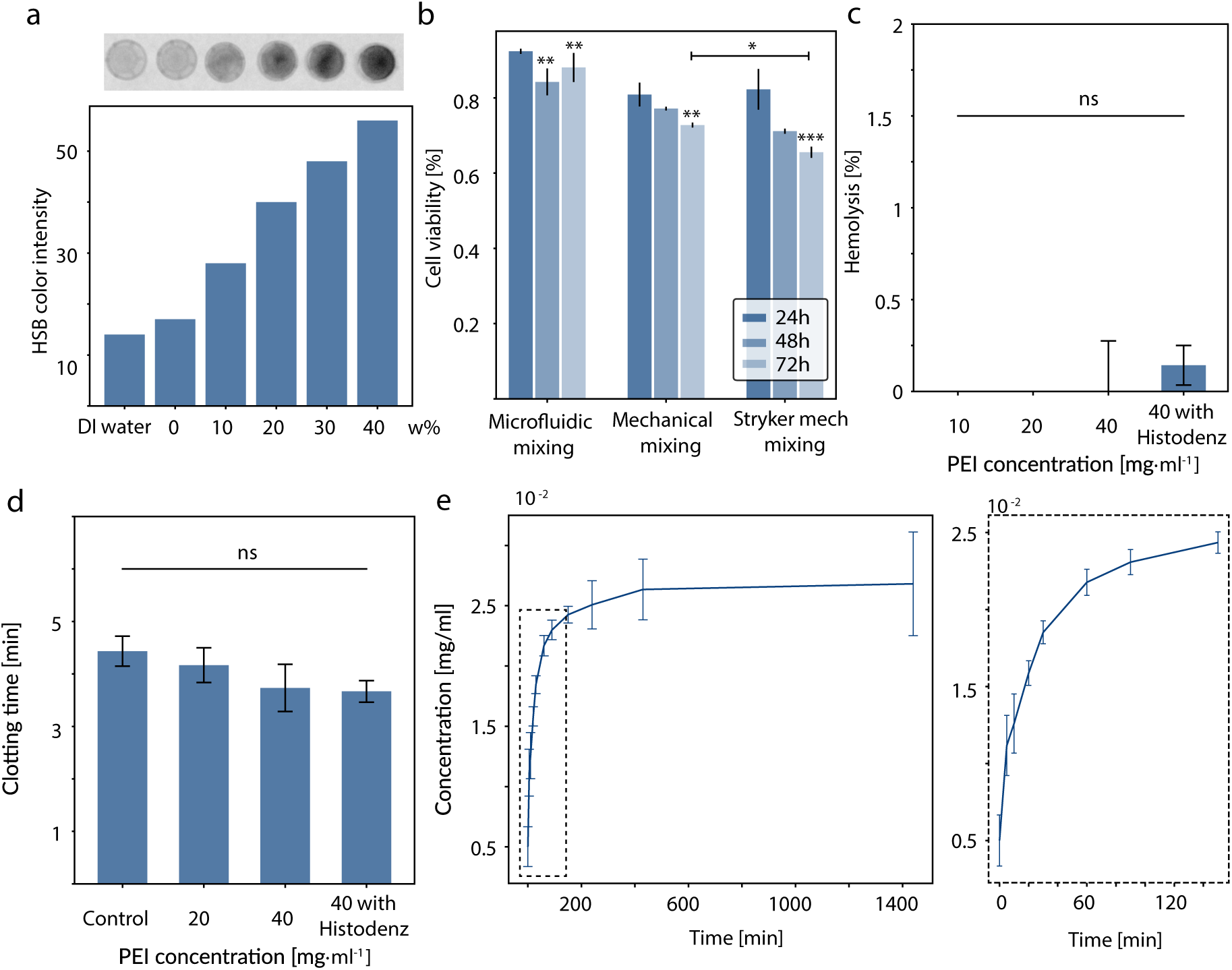
Biological and drug release characterization of PEI-PEG hydrogels containing 40% Histodenz. **a**, Fluoroscopy intensity of PEI-PEG hydrogels with Histodenz concentration ranging from 0 to 40 wt%. **b**, Cell viability of Human Umbilical Vein Endothelial cells (HUVEC) after incubating with PEI-PEG hydrogel for 24, 48, and 72 hours. Hydrogel mixing using microfluidics shows no toxicity over 72 hours. Mechanically mixed hydrogels, both our formulation and the Adherus AutoSpray formulation, show reduced cell viability over 24, 48, and 72 hours. **c**, Hemolysis values for different PEI concentrations and Histodenz loading suggest hemocompatibility of all materials. **d**, The clotting time is unaffected when incubated with porcine blood. Blood without additives is used as a negative control. **e**, Analysis of Doxorubicin release over 24 hours. Statistical significance was determined using one-way ANOVA. ns, not significant; \**p* < 0.05, \*\**p* < 0.01, and \*\*\**p* < 0.001.

### 2.4 Biocompatibility characteristics

The safety and effectiveness of PEI-PEG hydrogels must be evaluated in medical applications to demonstrate their potential as embolic materials. Key aspects of this evaluation include biocompatibility, thrombogenicity, and hemocompatibility. The biocompatibility of the hydrogel was tested against human umbilical vein endothelial cells (HUVEC) over a period of 72 hours (**Fig. 3b**). PEI-PEG hydrogel samples were incubated for 24, 48, and 72 hours, and cell viability was analyzed using an MTT assay.

The Adherus AutoSpray PEI-PEG formulation(Stryker) was used as an FDA-cleared reference. Both our formulation mixed in bulk and the commercial product showed an approximate 30% decrease in cell viability over 72 hours. In contrast, the hydrogel mixed using microfluidics exhibited almost no cytotoxicity and demonstrated a statistically significant increase in viability from 48 to 72 hours. These results indicate the improved biocompatibility of the PEI-PEG hydrogels generated using our microfluidic approach compared to those obtained using bulk methods and the reference commercial product. The compatibility with blood and associated hemolysis was assessed using a hemolysis assay (**Fig. 3c**). Hemolysis rates were all below 0.5% and were not detectable for lower PEI concentrations (< 20 mg/ml). Additionally, the presence of Histodenz also did not induce obvious hemolysis. Thrombus formation around an embolic material can improve stability and resistance to recanalization, so we evaluated thrombogenicity by measuring whole-blood clotting time. Wells coated with PEI-PEG hydrogel clotted in 3.5–4.0 min, whereas uncoated control wells clotted in 4.5 ± 0.3 min (**Fig. 3d and Supplementary Fig. 3**). This nominal 1 min difference was not statistically significant (p > 0.05); therefore, no meaningful decrease in clotting time was observed. We conclude that, under these in-vitro conditions, the hydrogel neither accelerates nor delays coagulation.

### 2.5 Drug release properties

To enhance the therapeutic potential of our hydrogels for chemoembolization, we evaluated their ability to load and release therapeutic agents. Specifically, we studied the drug release profile of PEI-PEG hydrogels using FDA-approved Doxorubicin (DOX) as a model drug.

For this evaluation, we prepared hydrogels in which DOX was pre-solubilized in the polymer (PEG) solutions before gel formation, resulting in a final DOX concentration of 5 mg/ml in the hydrogel. The release kinetics of DOX were tested over a period of 24 hours (**Fig. 3e**). The release profile showed that most of the DOX was released within the first 2 hours, indicating a burst release mechanism that rapidly achieves high local drug concentrations. Complete release was observed after 24 hours, demonstrating the hydrogel’s ability to deliver its drug payload efficiently within a short period. This rapid and complete release profile is beneficial for tumor treatment, which requires quick drug action^37^, making PEI-PEG hydrogels a promising candidate for chemoembolization therapies.

### 2.6 Microfluidic catheter and injectability

Having demonstrated the biocompatibility and drug delivery properties of the PEI-PEG hydrogel, and the ability to control its rheological properties using a microfluidic platform, we adapted this microfluidic design into a catheter system for endovascular interventions. The final design consists of a 1.5-meter-long multi-lumen tubing made from medical-grade polyurethane (Pellethane®), which is soft and suitable for intravascular use^38^. This material choice ensures biocompatibility and flexibility, making the device ideal for navigating the vascular system and precisely delivering the hydrogel. The tubing features four lumens configured for the individual injection of PEI, PEG, and PBS, which converge to a single-lumen microfluidic mixing chamber at the tip of the catheter. The tip is 1 cm-long, consistent with the previous microfluidic device design (**Supplementary Fig. 1a**). The Pellethane® tubing is additionally equipped with Pebax® jackets on the proximal end and magnetic elements on the distal end, to enhance pushability and navigability (**Supplementary Fig. 1d**). The Pebax® jackets are assembled along the length of the device with decreasing shore hardness, improving pushability through the complex neurovascular network and ensuring flexibility and navigation of the tip segment. NdFeB ring magnets are assembled on the distal tip segment, which is further processed using a thermal drawing process to decrease the diameter and local bending stiffness, thus improving magnetic navigation and actuation.

The catheter’s actuation performance was characterized using a Navion, an electromagnetic navigation system equipped with three parallel-configured electromagnets, and a Vicon motion tracking system. The catheter tip position was tracked at various lengths and distances from the electromagnetic navigation system. **Supplementary Fig. 1e** shows the catheter’s tip position in various anatomical locations, demonstrating its navigability and tip dexterity.

The delivery performance was evaluated by assessing injectability and block stability. Specifically, the injection force of the three liquids was measured using a microfluidic pump and an ATI Nano 17 force sensor. PEI and PEG were injected at a flow rate of 300 µL/min using a 1 mL syringe, followed by a 30-second pause, then PBS was injected at a flow rate of 1000 µL/min. The pause allows partial crosslinking within the mixing chamber, facilitating the measurement of the break-loose force. **Fig. 2i** shows the results, indicating a constant and reproducible injection force of less than 7 N for all injections. Additionally, the break-loose force of approximately 1.5 N ensures good reusability even after a short pause between injections.

The stability of the generated hydrogel block was assessed by injecting 0.28 mL of the hydrogel into a channel with a diameter of 3 mm, followed by a 15-minute setting time. The channel inlet pressure was then increased until reperfusion occurred. **Fig. 2j** illustrates the resulting pressure curve, showing the sudden failure and displacement of the embolic material. Although the failure mechanism varied between sudden displacement and partial recanalization, the pressure difference that caused failure was around 120 kPa (**Fig. 2k**), which is significantly higher than the pressure difference in blood vessels even for hypertensive individuals (≈ 25 kPa)^39^. This indicates that the PEI-PEG hydrogel is very stable in physiological conditions. As a control, we used the Adherus AutoSpray from Stryker and observed no statistically significant difference compared to our catheter system (**Fig. 2k**).

### 2.7 In vitro embolization in phantom models

To investigate the navigability and embolization performance in combination with the magnetic actuation of the catheter for precise hydrogel deployment, we performed in vitro experiments simulating the navigation in a mock human anatomy and embolization in clinical and simplified in vitro models. A full-body vascular model from Trandomed has been used to demonstrate the magnetic navigability under realistic flow conditions (**Supplementary Fig. 4b,c** and **Supplementary Video 2**). In this vascular model, we successfully navigated up to the M3 segment without using additional guidewires or other navigation devices. Additionally, we achieved embolization in the vertebral artery, providing promising results for embolization in high-flow anatomical structures (**Supplementary Fig. 4d** and **Supplementary Video 2**).

To further illustrate the versatility of our approach, custom anatomical models were fabricated out of silicone to reproduce realistic application scenarios of the embolic system. These anatomical models were either obtained from segmented clinical data or designed using CAD software, and focused on two key applications. Firstly, we demonstrated the application in an aneurysm filling while using a basilar aneurysm model (**Supplementary Fig. 5)**. The silicon model was perfused with physiological flow conditions, and the aneurysm was subsequently filled (**Fig. 4a** and **Supplementary Video 3**). We were able to track the filling under fluoroscopic imaging and observed a complete filling without perfusion within two injections (**Supplementary Fig. 5a,b)**. Secondly, navigation and embolization were demonstrated in a simplified liver model (**Supplementary Fig. 6**), in which we successfully embolized the left hepatic artery (**Fig. 4b** and **Supplementary Video 4**) and the feeding arteries to the VII Couninaud’s liver segment. All injections were trackable under fluoroscopic imaging (**Supplementary Fig. 6**), showing excellent visibility in all vessel sizes.

**Fig. 4.**
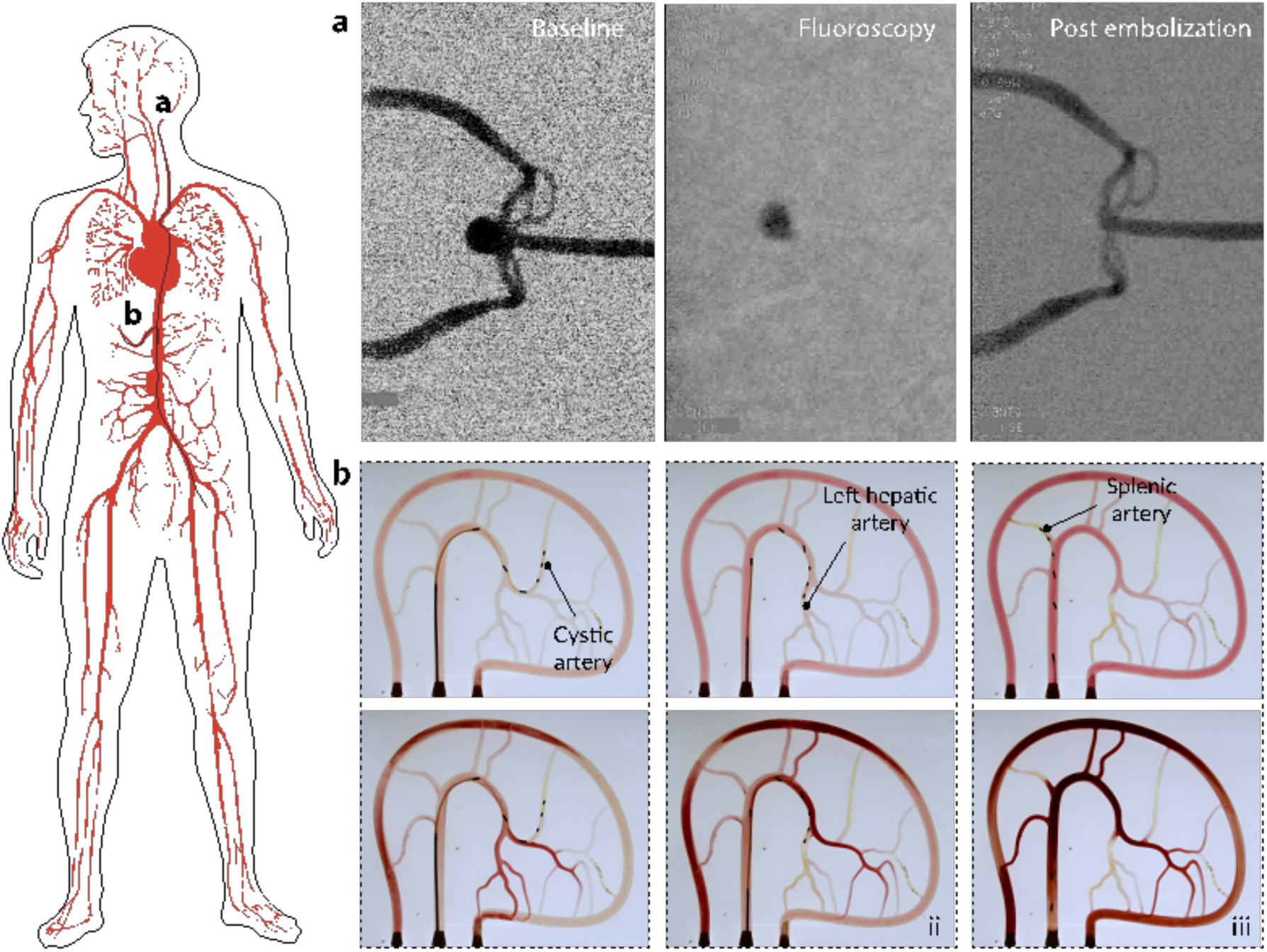
Transcatheter embolization (TCE) in in vitro models. (left) Schematic illustration of TCE applications in the human body, including the embolization of aneurysms and liver embolization with femoral catheter access. **a**, TCE in a basilar aneurysm model shows a complete filling without perfusion immediately after the injection. **b**, TCE in a liver model with multiple injections to targeted vessels and branches. Top images showing the navigation and catheter position while lower images are showing the perfusion of the model after each embolic material injection.

### 2.8 Ex vivo embolization in human placenta

Having demonstrated the validity and versatility of our approach using in vitro models, we extended our analysis by performing ex vivo experiments. We focused on human placentas, which are useful models for clinical training because their arteriovenous network closely resembling that of the human brain^40^. This similarity allows for training and tool validation in a realistic ex-vivo setup, accurately mimicking the conditions encountered in neuroradiology and surgery. Therefore, to further analyze the performance of our embolic device in a realistic human setting, we simulated multiple applications in a placenta. Specifically, we evaluated navigability in a perfused vessel system and validated the embolic hydrogel for embolization, chemoembolization, and hemorrhage treatment. The human placentas were first perfused with a heparinized saline solution according to the protocol reported by Julien Burel et al. ^41^ to remove the remaining blood and clear all vessels. An Avanti 5Fr introducer sheath was successfully placed in the umbilical vein, while a flat-tip steel needle was used to access the umbilical artery. This setup allowed complete perfusion of the placenta vessels, with an inflow in the umbilical vein and an outflow in the umbilical artery. The system was then connected to a peristaltic pump set up to precisely control the flow and pressure within the venous system.

#### 2.8.1 Navigability

The visibility and navigability of the developed microfluidic catheter under fluoroscopic guidance are crucial for a successful embolization procedure. To validate these aspects, we first performed catheterization of all vessels within a placental branch. The catheter was actuated using a mechanical advancer unit and controlled remotely via the electromagnetic Navion system from the control room. During the procedure, we tracked the catheter’s progress under fluoroscopic imaging to ensure accurate navigation. **Fig. 5a** and **Supplementary Video 5** illustrates the successful navigation through all vessels of the placental branch, demonstrating the catheter’s excellent visibility and navigability.

**Fig. 5.**
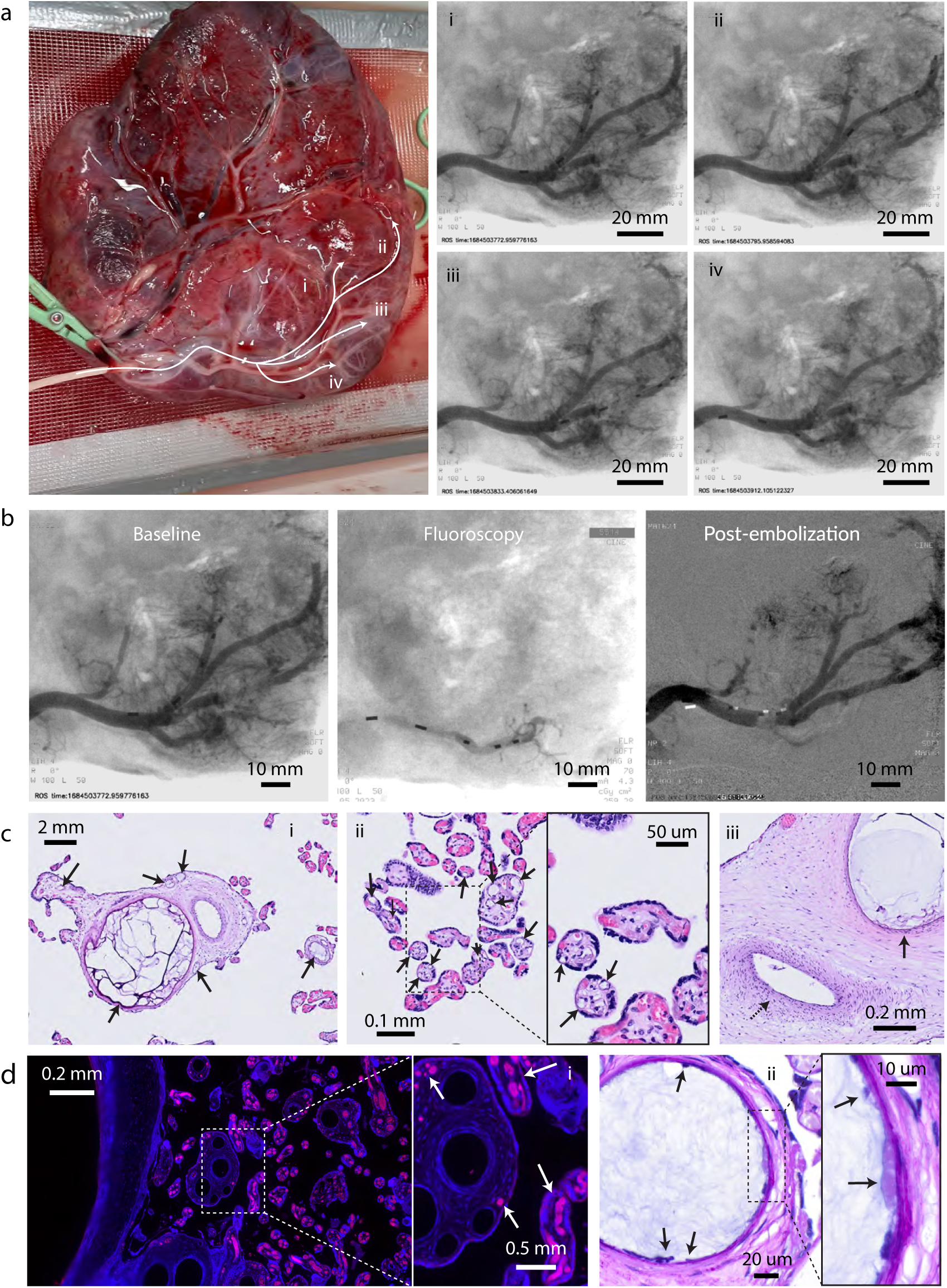
Microfluidic catheter navigation and selective embolization of sub-branches in an ex-vivo human placenta. **a**, A 4Fr introducer sheath is placed in the umbilical vein and used to access the right branch of the placenta. The microfluidic catheter was navigated under fluoroscopic imaging to the four main vessels i-iv of the targeted branch. **b**, Fluoroscopic imaging of the embolization process showing contrast medium injection, embolic material injection, and confirmation of complete embolization. **c**, Histopathological evaluation confirming complete filling in a wide range of vessel sizes, without visible tissue damage and cytotoxicity in combination with compression of tunica layers. **d**, Fluorescence histology evaluation of chemoembolic material illustrating the penetration of the chemoembolic material up to capillaries, with a fast release of DOX into the surrounding tissue in combination with a hydropic degeneration of the endothelial layer.

**Fig. 6.**
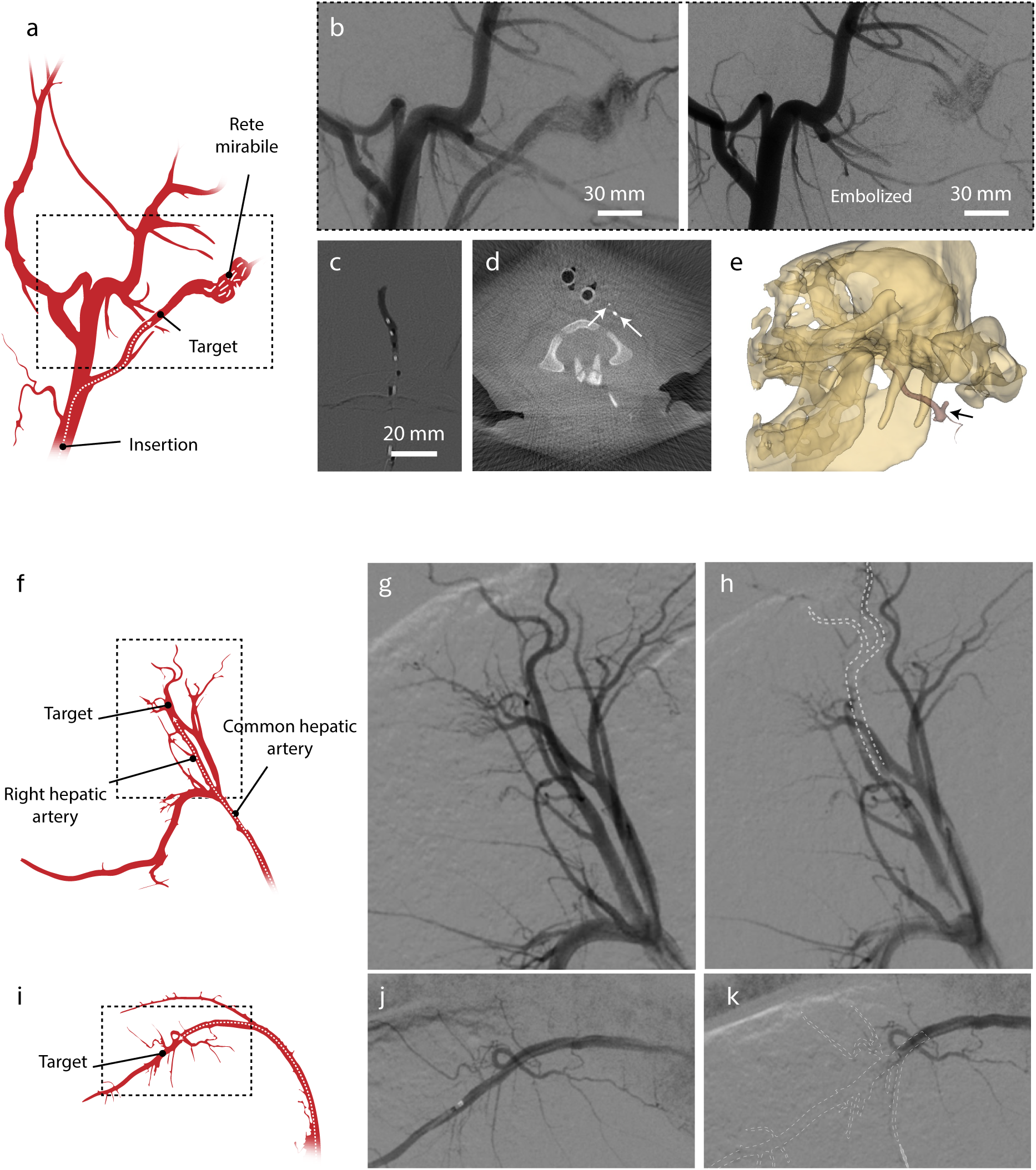
Selective embolization in a porcine model. **a**, Schematic of the cranial circulation showing navigation into the ascending pharyngeal artery (APA) supplying the rete mirabile. **b**, Digital-subtraction angiography (DSA) of the APA before (left) and after (right) embolization. **c**, Representative fluoroscopic. **d**, Post-procedural flat-panel CT (Xper CT). **e**, Three-dimensional reconstruction verifying precise intravascular localization of the embolic cast. **f**, Schematic of the right hepatic artery (RHA) target. **g**, DSA of the RHA before and **h**, after embolization. **i**, Schematic of the splenic artery (SA) target. **j**, DSA of the SA before and **k**, after embolization.

#### 2.8.2 Embolization

After demonstrating the catheter’s magnetic navigability in the human placenta, we subsequently evaluated its performance for embolization by targeting the occlusion of complete vascular branches. To achieve this, we selectively embolized sub-branches of the human placenta. The microfluidic catheter was positioned at the feeding vessel of a placental sub-branch, where the embolic material was injected at high flow rates to achieve complete filling up to the capillary structures. **Fig. 5b** and **Supplementary Video 6** illustrates the embolization process under fluoroscopic imaging. Initially, perfusion was assessed by injecting a contrast medium through the introducer sheath. Next, the embolic material was injected into the feeding vessel of the sub-branch while being monitored under fluoroscopic imaging. The injection was stopped once the embolic material had filled all targeted vessels and backflow around the catheter tip was observed. Finally, a second injection of contrast medium through the introducer sheath confirmed complete embolization, with no perfusion to the embolized vessels.

The complete filling up to smaller vessels was tracked under fluoroscopic imaging and confirmed by histopathology (**Fig. 5c i-ii**). We observed complete filling in all vessels ranging from 5 mm to 10 µm. Additionally, no acute signs of damage or inflammation were observable in the tunica intima and media. However, slight compression of the tunica layers was noted, visible as a thinner tunica layer and elongated shape of cells and nuclei in the filled vessels (**Fig. 5c iii**). This compression can be attributed to the marginal swelling of the embolic material.

#### 2.8.3 Hemorrhage

We also explored the potential of our embolic system for hemorrhage treatment. Embolization is an effective strategy for controlling bleeding by occluding the abnormal blood vessels responsible for the hemorrhage. To demonstrate this, we performed hemorrhage embolization by injecting a small amount of embolic material into a placental vessel previously punctured with a 14G needle. **Supplementary Fig. 7** and **Supplementary Video 7** illustrates the bleeding in the punctured vessel under physiological pressure. After embolization of the punctured vessel using the previously described procedure, we observed a slight extravasation of the injected material with an approximate volume of 1 µL. Reperfusion and bleeding were successfully inhibited up to an increased pressure of 250 mmHg. This demonstrates the potential of our embolic system to effectively treat hemorrhages by occluding the damaged vessels.

#### 2.8.4 Ex vivo chemoembolization in human placenta

Following the successful demonstration of the catheter’s navigability and embolization capabilities, we evaluated the application of the proposed technology for chemoembolization with localized drug delivery in a human placenta. For this purpose, we employed DOX as a model drug, loading it into a PEG solution. The microfluidic catheter was navigated to the feeding vessel of a placental sub-branch, where the DOX-loaded embolic material was then injected (**Supplementary Video 8**). The perfusion was maintained for 2 hours to ensure effective drug delivery before extracting samples for histopathological analysis.

Histological evaluation was performed to assess the penetration of the embolic material and the acute effect of DOX on the surrounding tissue. The results indicated complete penetration up to the capillaries and rapid release of DOX, as evidenced by the fluorescent signal in capillary-sized vessels and the increased fluorescent signal in the surrounding tissue (**Fig. 5d i**). Additionally, we observed an irregular endothelium with “bubbles” below the basal lamina (**Fig. 5d ii**). These “bubbles” are characteristic of the effects of chemotherapeutic agents and indicate hydropic degeneration, which involves increased water accumulation within cells. This phenomenon can be attributed to the decreased protective function of the endothelial layer, leading to compromised cellular integrity and increased water diffusion into the cells. The presence of hydropic degeneration suggests that DOX is effectively penetrating the targeted area and impacting the cellular structures, which is crucial for the therapeutic efficacy of chemoembolization.

### 2.9 In vivo embolization

In vivo models provide an essential platform for validating the proposed embolization process in a setting that closely mimics the actual field of application. This similarity allows for validation in models that express realistic blood perfusion, pulsatility, fluoroscopic contrast, complex vascular structures, and dynamic vascular deformations. To this end, we evaluated the embolization performance and material visibility in a live porcine model.

Selective embolization was assessed in a porcine model across three vascular areas, the ascending pharyngeal artery (APA) supplying the rete mirabile, the right hepatic artery (RHA), and the splenic artery (SA), using our soft-robotic microfluidic catheter. Vascular access was obtained with a 7 F femoral sheath in the left femoral artery and a 7F guide catheter was used to explore the coeliac trunk. Fresh precursor solutions (PEG 332 mg/mL; PEI 20 mg/mL) were prepared immediately before each procedure and co-infused at equal flow rates of 300 µL/min. Infusion continued until angiography showed complete stasis in the target segment. The two streams mixed at the catheter tip and gelled within seconds, consistent with in-vitro and ex-vivo results.

**Fig. 6a** schematizes the cranial circulation and the APA rete mirabile pathway. **Fig. 6b** shows digital-subtraction angiography (DSA) before (left) and after (right) embolization, demonstrating accurate, segment-selective occlusion with preservation of adjacent branches (**Supplementary Video 9**). A representative fluoroscopic frame during delivery (**Fig. 6c**) illustrates the hydrogel’s intrinsic radiopacity and a smooth intraluminal cast. Immediately post-embolization, flat-panel CT (Xper CT) confirmed a sharply confined, high-attenuation cast in the intended segment (**Fig. 6d**), and the 3-D reconstruction verified precise intravascular localization (**Fig. 6e**).

**Fig. 6f** provides the RHA scheme and **Fig. 6g,h** presents the paired DSA images before (**Fig. 6g**) and after (**Fig. 6h**) embolization, documenting complete and selective occlusion without reflux or distal migration (**Supplementary Video 10**). The SA is schematized in **Fig. 6i**, with the corresponding DSA pair before (**Fig. 6j**) and after (**Fig. 6k**) embolization showing an equivalent instant and stable occlusion of the targeted segment (**Supplementary Video 11**). Post-procedural flat-panel CT for both visceral areas corroborated intravascular confinement and cast stability; the complete CT series is provided in the Supplementary Information.

No complications such as vasospasm or non-target embolization were observed during the interventions. Post-operative CT imaging confirmed the stability of the embolic material, supporting its suitability for high-flow vascular environments (**Supplementary Fig. 13**). Importantly, no adverse reactions such as inflammation were observed. Post-procedural histopathological analysis further confirmed the absence of any negative tissue responses, indicating the biocompatibility and safety of the hydrogel embolic material across all tested sites (**Supplementary Fig. 14 and Supplementary Fig. 15**).

### 2.10 In vivo chemoembolization in rat models

After demonstrating targeted drug delivery for ex-vivo chemoembolization experiments performed in the human placenta and in vivo embolization in a porcine model, we analyzed the potential of the DOX-loaded embolic hydrogel using in vivo rat tumoral-liver models. First, we assessed the inhibitory effects of the DOX-loaded hydrogels on luciferase-labeled rat liver cancer cells (CBRH-7919-luc). We specifically used this labeling strategy to further improve tumor analysis in the post in vivo animal models. Hence, we incubated the labeled cancer cells with samples of the embolic hydrogel containing different concentrations of DOX for 48 hours. After this period, cell viability was significantly reduced in a DOX concentration-dependent manner, as determined by the MTT assay, indicating effective cytotoxic activity because of the drug delivery (**Supplementary Fig. 8**). Notably, the hydrogel without drug exhibited excellent biocompatibility, confirming our previous in vitro observations. Subsequently, a rat model with orthotopic liver tumors was established using these luciferase-labeled rat liver cancer cells. The study protocol is detailed in **Fig. 7a**, which illustrates the timeline from tumor implantation 10 days before the treatment day (Day 10) to final analyses conducted 12 days after the treatment (Day 12). Initial in vivo bioluminescence imaging on Day-10 ensured uniform tumor load across the experimental groups, confirming that all rats exhibited comparable tumor presence before the chemoembolization (**Fig. 7b**). Additionally, quantitative analysis verified no significant differences in the initial tumor load, establishing a consistent baseline for assessing the efficacy of the chemoembolization treatment (**Fig. 7c**).

**Fig. 7.**
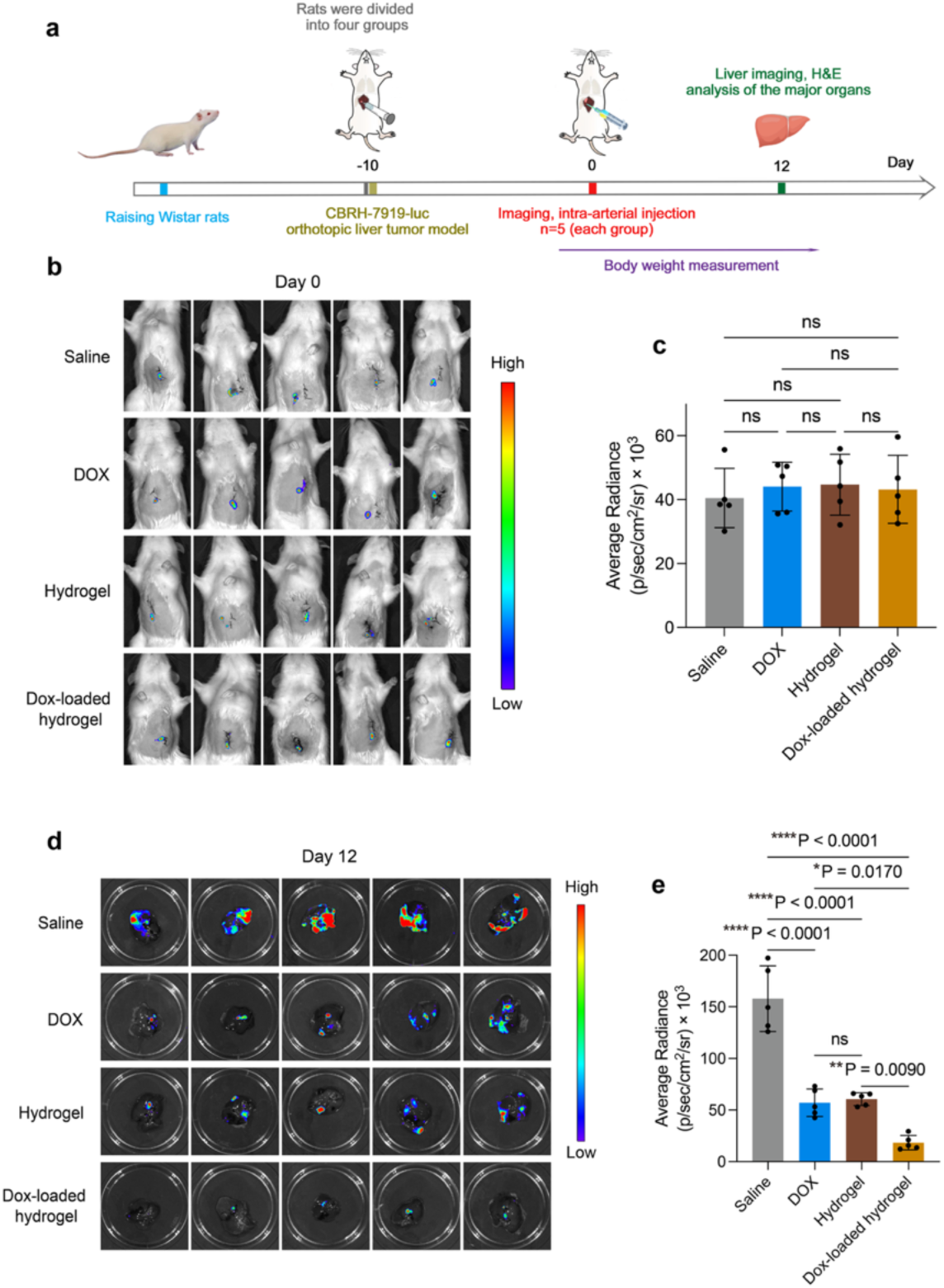
Efficacy of Hepatic Artery Chemoembolization in Rats with CBRH-7919-luc Hepatic Tumors. **a**, Schematic of the experimental design implementing our theranostic approach. **b**, Representative in vivo bioluminescence images of CBRH-7919-luc tumors in the liver before intra-arterial hydrogel delivery on Day 0, demonstrating tumor presence. **c**, Quantification of bioluminescent radiance from liver pre-treatment, averaged across n = 5 rats (Data presented as mean ± SEM). **d**, In vivo bioluminescence imaging post-treatment, and **e**, Corresponding quantified bioluminescent radiance of the isolated liver on Day 12 after various treatments (n = 5, data presented as mean ± SEM). Statistical analysis was performed using one-way ANOVA with a Tukey post-hoc test (Fig. 7c**,e**). Significance levels are indicated: *p < 0.05, **p < 0.01, ****p < 0.0001 versus control.

Following baseline assessment, twenty rats were randomized into four groups (n = 5) depending on the treatment administered: control (Saline), DOX, hydrogel, and DOX-loaded hydrogel. The DOX and saline were administered via tail injection, while the hydrogel and DOX-loaded hydrogel were delivered directly to the liver tumors by embolizing the main tumor feeding artery. Importantly, the same dose of DOX was used for both the tail injection and hydrogel delivery. Post-treatment analysis, conducted through in vivo bioluminescence imaging 12 days after the intervention, revealed that the rats treated with saline continued to show, as expected, an increase in tumor size. In contrast, the groups treated with DOX or hydrogel (DOX-free) exhibited similar impaired tumor growth with tumor luminescence that was approximately one third of that observed in the saline group (**Fig. 7e**). This demonstrated that merely restricting nutrient supply to the tumor through embolization had a similar effect to the pure chemotherapeutic action. More importantly, the group treated with the DOX-loaded hydrogel exhibited a substantially lower tumor growth, with tumor luminescence intensity being 32% of that observed in the group treated with DOX alone (**Fig. 7e**). This indicated that the DOX-loaded hydrogel reduced tumor viability, and enhanced tumor targeting and treatment efficacy (**Fig. 7d**). Further histological examination of liver sections stained with Hematoxylin and Eosin (H&E) supports the bioluminescence findings. The DOX-loaded hydrogel group showed minimal tumor infiltration compared to the other groups (**Supplementary Fig. 9a**). This demonstrates the synergistic effect of doxorubicin’s chemo cytotoxicity and the embolic properties of the hydrogel. Chemoembolization performed using our embolic formulation and microfluidic catheter not only blocks the nutrient supply to the tumor but also ensures controlled and targeted delivery of doxorubicin directly into the tumor, enhancing the overall therapeutic efficacy.

Histological analysis also confirmed the absence of significant damage in major organs, affirming the safety and targeted efficacy of the hydrogel formulation. Additionally, throughout the study, the health and body weight of the rats were monitored to assess treatment tolerability. No significant adverse effects were observed, indicating that the interventions were well-tolerated by all groups (**Supplementary Fig. 9b**).

## 3 Discussion

In this study, we introduced a novel two-component hydrogel embolic material composed of activated PEG and PEI, delivered through a custom-designed microfluidic catheter system. The hydrogel demonstrated immediate crosslinking upon mixing, tunable mechanical properties, excellent biocompatibility, enhanced visibility under fluoroscopic imaging, and effective drug-loading capabilities, addressing several limitations of current embolization agents.

Rheological characterization revealed that the PEI-PEG hydrogel exhibits solid-like behaviour ideal for embolization applications. Its shear-thinning and thixotropic properties facilitate catheter delivery by allowing it to flow under shear stress during injection and recover its viscosity once in place, enhancing embolus stability. The significant improvement in mechanical properties observed with microfluidic mixing compared to bulk mixing suggests that controlled mixing at the microscale promotes efficient and homogeneous crosslinking. This results in a hydrogel with higher stiffness and better performance. Moreover, the ability to tune the hydrogel mechanical properties by adjusting injection flow rates and PEI concentrations offers adaptability to various vascular conditions, making the hydrogel promising for precision medicine approaches and for different clinical scenarios.

Incorporating Histodenz as a contrast agent effectively increased the hydrogel radiopacity, allowing for excellent visibility under fluoroscopic imaging. In addition, biocompatibility assessments showed that the hydrogel mixed using microfluidics exhibits minimal cytotoxicity toward human endothelial cells, outperforming bulk-mixed formulations. Hemolysis assays confirmed hemocompatibility, with hemolysis rates well below acceptable thresholds, indicating that the hydrogel is safe for contact with blood.

The incorporation of magnetic elements into the microfluidic catheter facilitated precise navigability and positioning of the hydrogel, improving flexibility, and allowing navigation through complex vascular networks without the need for additional guidewires. The catheter’s excellent navigability was demonstrated in vitro, ex vivo in human placenta, and in vivo in porcine models. Ex vivo experiments in human placenta demonstrated the hydrogel’s effectiveness in embolization, hemorrhage control, and chemoembolization, with histopathological analyses confirming complete vessel occlusion up to the capillary level without inducing acute tissue damage or inflammation, indicating safety and efficacy. In vivo studies further validated these findings, with successful embolization in porcine spleen, liver, and brain without complications like vasospasm or non-target embolization.

The hydrogel’s capacity for drug loading and controlled release was demonstrated using doxorubicin as a model chemotherapeutic agent. The release profile showed rapid drug delivery within the first few hours, beneficial for achieving high local concentrations immediately post-embolization. This enhances therapeutic efficacy while potentially minimizing systemic exposure and side effects. The successful chemoembolization observed in the rat liver tumor model underscores the hydrogel’s potential for targeted cancer therapy in clinical applications, by effectively combining mechanical occlusion with localized chemotherapy.

In summary, the PEI-PEG hydrogel embolic material and magnetically navigable microfluidic catheter offer a versatile and effective solution for various clinical applications, as demonstrated in ex vivo and in vivo models. This technology could significantly improve endovascular interventions, offering a new paradigm in treating vascular diseases and tumors.

## 4 Methods

*Rheology*: PEI–PEG hydrogel variants were prepared and subjected to rheological characterization across relevant operating windows, with viscoelastic responses compared across multiple formulations. Rheology measurements have been carried out on an Anton Paar MCR 502 rotational rheometer. The results were gathered with an aluminum plate-plate, sand-blasted geometry of 25 mm in diameter, as it ensured that no slip could take place. A temperature of 37°C was kept throughout the experiments to simulate the environment of the human body. The material analysis included frequency and amplitude sweeps, thixotropy tests and crosslinking tests. The frequency sweep tests covered 7 points per decade, with frequencies ranging from 0.1 rad s^-1^ to 100 rad s^-1^ at a constant strain amplitude of 1%. The amplitude sweep recorded 7 points per decade, with strain amplitudes spanning between 0.1% and 1000%, at a constant angular frequency of 1 rad s^-1^. Thixotropy tests were conducted at 10 rad s^-1^ under strain oscillation between 0.1% (low strain) for 2 minutes and 100% (high strain) for 2 minutes to examine the recoverability of PEI-PEG. 5 cycles were run on each sample to observe any prevailing trends. Finally, the crosslinking tests consisted of running the rheometer at a constant frequency of 10 rad s^-1^ and strain amplitude of 1%, measuring every 3 seconds. To ensure that all the time-dependent experiments had a common starting point, a chronometer was used to measure 1 minute after the hydrogel had been deposited on the plate, only then starting the test. Apart from the comparison between mixing strategies, PEI-PEG hydrogels were deposited onto the rheometer plate via injection through the microfluidic catheter. The mechanically mixed hydrogel was pipetted onto the plate and mixed with the pipet tip.

*Numerical simulations:* The flow and mass transport inside the multi-lumen microfluidic catheter device were simulated using computational fluid dynamics. The governing equations for these simulations are the Navier-Stokes equation for incompressible flow, the continuity equation, and the species transport equation, which are respectively given by:

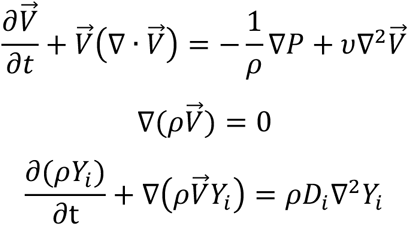

where 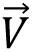 is the velocity vector, *ρ* is the density, *P* is the pressure, *ν* is the kinematic viscosity, *Y*_i_ is the mass fraction of species i and *D*_i_ is the diffusion coefficient of species i. These equations were solved numerically using the finite volume method with second-order upwind discretization. A steady-state double precision pressure-based solver was used for solving the equations, with a coupled algorithm for velocity-pressure coupling. Convergence was assumed to be achieved when residuals were lower than 10^-6^, given that stricter criteria produced similar concentration and velocity fields.

The boundary conditions used in the simulations were chosen to mimic experiments. Specifically, PEI and PEG solutions were introduced at each of the central lumens, while water (PBS) was introduced at the external lumens, typically at a flow rate of 300 µL/min each (total flow rate of 900 µL/min). The properties of the fluids used in simulations were assumed to be those of water, based on experimental confirmation that their properties are similar (density = 1000 kg/m^3^, viscosity = 0.001 Pa·s). Furthermore, the diffusion coefficients of PEI and PEG were considered to be 1.9 x 10^-10^ m^2^/s based on an empirical equation that relates the diffusion coefficient with the molecular weight of PEG molecules.^42^ We assumed that the reaction-diffusion (RD) zone where most of the hydrogel forms is the region of the microfluidic device where the concentrations of PEI and PEG exceed 1 % of their inlet concentration simultaneously. No-slip conditions were assumed at the device walls, and atmospheric pressure was assumed at the outlet.

*Mesh testing and validation of numerical simulations*: We performed preliminary simulations of the flow and mass transport of PEI and PEG in the multi-lumen catheter to identify the coarsest mesh that could produce mesh-independent results. We found that a mesh containing approximately 10 million cells produced results like those obtained using a much finer mesh with approximately 14 million cells. Specifically, the velocity profile along the radius of the device outlet and the RD zone at the outlet, i.e. the region in which the concentration of PEI and PEG exceed 1 % of their initial concentration, were similar when using both meshes (**Supplementary Fig. 10a,b**). Therefore, we chose the mesh containing 10 million cells to use in all simulations. The velocity profile along the radius of the device outlet was compared with the theoretical velocity profile in a circular pipe and complete agreement was found (**Supplementary Fig. 10a**),^43^ showing that our simulations can accurately capture the relevant transport phenomena.

*Swelling*: Swelling of the embolic material is a highly relevant characteristic in an embolization procedure, as it dictates the long-term behavior of the hydrogel in a physiological environment. It is desirable for the mass swelling to be low for avoiding any significant increase in arterial pressure and potential rupture or recanalization. The method employed for characterizing the swelling behavior of PEI-PEG hydrogels followed that outlined by Modak et. Al..^44^ This consisted of measuring the change in mass for a set of samples of a known geometry over time. Hydrogel discs with a diameter of 15 mm and a height of 5 mm were prepared. The disks were immersed in PBS and incubated at 37°C. The weight change was measured on a scale after removing the excess humidity with absorbing paper.

*Injectability*: Injectability was assessed by measuring the force applied by the microfluidic pump onto the syringe piston. A Nano 17 6-axis force sensor (ATI industrial automation) was mounted on the microfluidic pump and used to monitor the injection forces. Injection forces of PEI and PEG were measured on a 130 cm long microfluidic catheter with an injection rate of 300 µL/min. The injection force of PBS is not only an indicator of injection pressure but also of break-loose force of the Hydrogel in the mixing chamber at the catheter tip. After the injection of PEI and PEG, all flows were turned off for 10 s, and then PBS was injected at a flow rate of 1000 µL/min while monitoring the applied force.

*Block stability*: The stability of the injected material under pressure is an important parameter to assess the stability and migration of the embolization material. To assess the stability against pressure, we injected a total volume of 0.28 mL of PEI-PEG hydrogel into a channel with a diameter of 3 mm. To assess the long-term stability, we waited for 30 min before applying pressure. Subsequently, the pressure was gradually increased while monitoring the pressure with a Honeywell 19C015PG5K pressure sensor. As a reference, we also analyzed the pressure resistance of the same volume of the Stryker hydrogel.

*Cell Culture:* Human Umbilical Vein Endothelial Cells (HUVECs) were cultivated under controlled conditions in an incubator set to maintain a 5% CO_2_ atmosphere at a constant temperature of 37°C. The culture medium used was Dulbecco’s Modified Eagle Medium (DMEM, Gibco), enhanced with a 12% Fetal Bovine Serum (FBS) supplement, in addition to 1% penicillin/streptomycin for antimicrobial protection. To ensure optimal cell growth and maintenance, the medium was replenished every three days, with the cells undergoing passaging procedures biweekly. The rat hepatoma cell line, CBRH-7919-luc, supplied by Shanghai Zhong Qiao Xin Zhou Biotechnology Co., Ltd., was cultured in Dulbecco’s Modified Eagle Medium (DMEM) enriched with 10% fetal bovine serum (both from Gibco), 100 IU/mL penicillin G, and 100 μg/mL streptomycin. Cultures were maintained under a humidified atmosphere of 5% CO2 and 95% air. Prior to implantation, cell viability was assessed using trypan blue exclusion, ensuring a viability exceeding 90% for each tumor implantation procedure.

*Cell viability*: Hydrogel samples were subjected to sterilization using UV light for a duration of 2 h. Human Umbilical Vein Endothelial Cells (HUVECs) were then cultivated in 24-well culture plates at a density of 10,000 cells per well in 1 mL Dulbecco’s Modified Eagle Medium (DMEM) for 24 h. These cells were incubated in the presence of the sterilized hydrogel samples for time periods of 24, 48, and 72 h, respectively. Following the incubation period, 50 μL of sterilized MTT solution (3 mg/mL in Phosphate-Buffered Saline, sourced from Thermo Fisher Scientific) was introduced to each well. After subsequent incubation at 37 C° for 4 h, the medium was carefully removed from each well. To dissolve any residual insoluble formazan crystals, 1 mL of Dimethyl Sulfoxide (DMSO) was added. Finally, the absorbance was quantified at a wavelength of 560 nm utilizing an Infinite® 200 PRO microplate reader from TECAN. Cell viability was then determined and expressed as a percentage in relation to the untreated control cells.

*Visibility*: Imaging of the embolic material was achieved by fluoroscopic imaging using a Ziehm Vision FD (Ziehm Imaging). The fluoroscope was used for visibility assessment and in vitro and ex-vivo validation. A mono-plane angiography system (Allura Xper FD20 fluoroscope, Philips N.V., Amsterdam, Netherlands) was instead used for in vivo experiments.

*Hemolysis*: The hemocompatibility of the composite hydrogels was assessed via interaction with healthy porcine blood procured from the slaughterhouse (SBZ Schlachtbetrieb Zürich AG). The freshly obtained red blood cells were subjected to centrifugation at 500x g for 5 min. This was followed by successive washes using a Phosphate Buffered Saline (PBS) solution, maintained at a pH of 7.4. Concurrently, composite hydrogel samples were conditioned at 37 °C in a PBS solution, and the procured blood samples were diluted using the same PBS solution. For a duration of one hour, these blood samples were incubated at a steady temperature of 37 °C. Control samples, both positive and negative, were similarly incubated at 37 °C. The positive control was subjected to a 1% Triton solution (Sigma Aldrich), while the negative control was maintained in the PBS media. Following incubation, samples were centrifuged at 1600 g for a duration of 5 min. The supernatant’s absorbance was then measured at a wavelength (λ) of 540 nm using a microplate reader, providing an estimate of the hemoglobin released during the process. The hemolysis percentage was subsequently calculated using the prescribed equation.

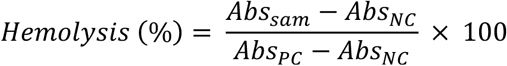

*Clotting time*: We utilized 100 μL hydrogel samples to coat the base of sequential wells within a 96-well plate. We subsequently prepared a solution comprising citrated rat blood and calcium chloride (CaCl_2_) in a 10:1 ratio, subjecting the mixture to a brief vortexing period of 10 seconds. Following this, a 50 μL aliquot of this blood sample was meticulously dispensed into the prepared wells. A series of predetermined time points were then identified (at intervals of 1, 2, 3, 4, 5, and 6 min). At each of these junctures, each well underwent a thorough washing procedure with PBS to completely remove all uncoagulated blood components. The duration required for the formation of a solid blood clot within each well was accurately documented, and this time frame was subsequently denoted as the blood clotting time.

*Doxorubicin release*: The capabilities of PEI-PEG hydrogels to carry and release DOX have been characterized by its diffusion profile. Hydrogel samples with a volume of 40 µL and a drug concentration of 5 mg/mL were prepared and immersed in an Eppendorf tube with 1 mL of PBS at 37°C. The tubes were placed in a thermomixer at 37°C and 500 rpm. At given time intervals, 20 µL of solution was removed and diluted in 150 µL of PBS solution. The fluorescent intensity at 596 nm with an excitation at 494 nm was analyzed with a well plate reader (Tecan Microplate Reader). Measurements were run until a steady state was reached. The fluorescence-to-concentration calibration curve was obtained previously by preparing a series of DOX dilutions in PBS.

*In vitro model fabrication:* Silicone models have been manufactured for in vitro testing of the catheter navigation and delivery of the PEI-PEG hydrogel. These models were produced by casting silicon on a previously 3D-printed (FDM printer) ABS geometry. Once molded, the ABS structure could be dissolved in acetone. Simplified 2D geometries were drawn in Onshape Inc., while more complex anatomical models were generated based on clinical data. Barb connectors were used to connect the model to a pump setup providing a physiological flow during the experiments.

*Magnetic manipulation*: Magnetic manipulation was achieved using a Navion, an electromagnetic navigation system generating magnetic fields up to 20 mT within a 25 cm side cube in front of the coil’s surface.

*Robotic advancer unit*: Advancement and retraction of the catheters were achieved with a robotic advancer unit consisting of a 3D printed casing that enables insertion and removal of the catheters. The advancer is controlled with a Maxon EPOS microcontroller (EPOS4 Compact 24/1.5 EtherCAT, Maxon Motors), providing control of the advancer’s position and velocity.

*Placenta preparation*: Human placentas were obtained from patients with written consent and approval from the Ethical Committee of the District of Zürich (study Stv22/2006). The placentas were first perfused with a heparinized saline solution according to the protocol reported by Burel et al. ^41^. Once all vessels were free of blood, an Avanti 5Fr introducer sheath was placed in the umbilical vein while a flat-tip steel needle was used to access the umbilical artery. This allows complete perfusion of the placenta vessels with an inflow in the umbilical vein and an outflow in the umbilical artery. The system was then connected to a peristaltic pump setup consisting of multiple stepper motor-driven peristaltic pumps in order to precisely control the flow and pressure within the venous system. Flow conditions were assessed with a DSA technique. The venous pressure was instead monitored with a pressure gauge and controlled within physiological conditions. The placenta was perfused with a saline solution throughout the experimental procedure.

*Histopathology*: Several different locations of the placenta were fixed in 4% phosphate buffered formalin for 24h. Then, they were dehydrated in an ascending alcohol series and embedded in paraffin. The blocks were cut in 2-3 µm sections and stained with routine H&E protocol. For fluorescence imaging, the slides were stained with DAPI (1: 1000 for 5min, Thermo fisher scientific, Basel, Switzerland). For analysis on H&E as well as the autofluorescence was used the Olympus slide scanner VS200, (Evident Technology Center Europe GmbH, Berlin, Germany).

*In vivo porcine model:* A 4-month-old, 50 kg female domestic pig of Swiss large white breed was used for *in vivo* validation under general anesthesia. The vital parameters of the pig were monitored by a veterinarian team throughout the procedure, and the pig was euthanized after the procedure. The animal study was approved by the local Committee for Animal Experimental Research (Cantonal Veterinary Office Zurich, Switzerland) under license number ZH213/2019. The ACT varied between 150 and 220.

A mono-plane Allura X-per angiography system (Philips, The Netherlands) was used. Arterial access was obtained by using a 7F sheath into the left common femoral artery (CFA). Under fluoroscopic guidance, exploration of the celiac trunk with a 7F guide catheter (7F Envoy MPC, Cerenovus) and a 0.035-inch standard guide wire (Bentson Wire Guide). The guide catheter was continuously flushed with heparinized saline (NaCl 0.9%, bbraun, Germany). As contrast agent Ultravist 300 (Iopromidum 300mg/ml, Bayer, Germany) was used. After diagnostic angiographic runs of the branches of the celiac run, the distal segment of the splenic artery (SA) was explored manually using the specifically designed embolization catheter under fluoroscopic guidance using smart map, positioning of the distal catheter tip at the proximal distal SA segment. The embolization was performed under fluoroscopic control using glue map. After the embolization, the catheter was withdrawn, and control angio-runs were performed. Then, the common hepatic artery (CHA) was explored using the same 7F guide catheter. After a diagnostic angiographic run of the branches of the CHA, the embolization catheter was manually advanced into the trunk of the right hepatic artery (RHA). After positioning of the distal tip of the catheter into the RHA, the embolization was performed under fluoroscopic control using a glue map. After the embolization, the catheter was withdrawn and control angiographic runs were performed showing the complete occlusion of the distal branches of the RHA. Similarly, selective embolization of the ascending pharyngeal artery was also performed. A thoraco-abdominal computed tomography (CT) with portal-venous and arterial phases were obtained from the pig. Finally, the pig was euthanized and the liver and spleen were obtained for histological analysis. Imaging analysis was performed OsiriXMD v14.1.1.

### In vivo chemoembolization model

Animal Model and Tumor Implantation: In the CBRH-7919 hepatoma model, Wistar rats weighing 180-220 g (procured from Beijing HFK Bioscience Co., Ltd.) were anesthetized with sodium pentobarbital. Subsequently, a surgical procedure was performed to expose the left medial lobe of the liver, into which 0.1 mL of the CBRH-7919 hepatoma cell suspension (1 × 10^8^ cells/mL) was directly injected under the hepatic capsule. The incision was then sutured closed. On day 10 post-implantation (designated as Day 0 for subsequent interventions), in vivo bioluminescence imaging was employed to confirm the absence of significant differences in tumor development among the various rat groups.

Therapeutic Intervention and Grouping: Twenty orthotopic liver tumor-bearing rats were randomly divided into four experimental groups: Saline, Doxorubicin (DOX), Hydrogel, and Dox-loaded Hydrogel, with five rats per group. The celiac, hepatic, and gastroduodenal arteries were meticulously isolated. Ligatures were placed on the gastroduodenal artery both distally and proximally to the intended puncture site, with an additional temporary ligature on the celiac artery to halt arterial flow. Utilizing a double-syringe system (Mixpac, medmix Switzerland AG) equipped with a custom mixing needle, the gastroduodenal artery was injected the hydrogel loaded with DOX (100 μL, containing 1 mg DOX, 16.6 mg PEG, and 1 mg PEI) was administered towards the hepatic artery. Following the injection, the proximal part of the gastroduodenal artery was ligated, the celiac artery ligature was removed, and hepatic arterial flow was reinstated.

Post-Treatment Analysis and Histological Examination: On day 12, the rats were euthanized, and major organs, including the liver, were harvested. In vivo bioluminescence imaging was reiterated to monitor tumor progression post-treatment. Subsequently, the organs were fixed in formalin and subjected to hematoxylin and eosin (H&E) staining. Additionally, body weights of the rats were monitored bi-daily over the 12-day period, aiding in the assessment of treatment tolerability.

## Supporting Information

Supporting Information is available from the Wiley Online Library or from the author.

## Acknowledgements

This work was supported by the ITC-InnoHK grant 16312 and SNSF for the FLUENS project (No. 10005033). Part of the work was supported by the European Union’s H2020 research and innovation programme under Grant Agreements (No. 952152). TSM and JPV acknowledge the support by LA/P/0045/2020 (ALiCE), UID/00532/2025 and UID/PRR/00532/2025 (CEFT), funded by Portugal through FCT/MECI.

## References

1. Wáng, Y.-X. J., De Baere, T., Idée, J.-M. & Ballet, S. Transcatheter embolization therapy in liver cancer: an update of clinical evidences. Chin J Cancer Res 27, 96–121 (2015).

2. Yonemitsu, T. et al. Evaluation of transcatheter arterial embolization with gelatin sponge particles, microcoils, and n-butyl cyanoacrylate for acute arterial bleeding in a coagulopathic condition. J Vasc Interv Radiol 20, 1176–1187 (2009).

3. Fiore, F. et al. Transarterial embolization (TAE) is equally effective and slightly safer than transarterial chemoembolization (TACE) to manage liver metastases in neuroendocrine tumors. Endocrine 47, 177–182 (2014).

4. Jiang, H. et al. Antiangiogenic therapy enhances the efficacy of transcatheter arterial embolization for hepatocellular carcinomas. Int J Cancer 121, 416–424 (2007).

5. Hu, J. et al. Advances in Biomaterials and Technologies for Vascular Embolization. Advanced Materials 31, 1901071 (2019).

6. Do, J. et al. Controlled release of doxorubicin from bio-resolvable drug-eluting bead of GelMA-SPMA graft copolymer for embolotherapy of liver cancer. Journal of Drug Delivery Science and Technology 105, 106601 (2025).

7. Lewis, A. L. & Dreher, M. R. Locoregional Drug Delivery Using Image-guided Intra-arterial Drug Eluting Bead Therapy. J Control Release 161, 338–350 (2012).

8. Kettenbach, J. et al. Drug-loaded microspheres for the treatment of liver cancer: review of current results. Cardiovasc Intervent Radiol 31, 468–476 (2008).

9. Hu, J. et al. Advances in Biomaterials and Technologies for Vascular Embolization. Advanced Materials 31, 1901071 (2019).

10. Young, S., Rostambeigi, N. & Golzarian, J. The Common but Complicated Tool: Review of Embolic Materials for the Interventional Radiologist. Semin Intervent Radiol 38, 535–541 (2021).

11. Pal, A., Blanzy, J., Gómez, K. J. R., Preul, M. C. & Vernon, B. L. Liquid Embolic Agents for Endovascular Embolization: A Review. Gels 9, 378 (2023).

12. Jordan, O., Doelker, E. & Rüfenacht, D. A. Biomaterials Used in Injectable Implants (Liquid Embolics) for Percutaneous Filling of Vascular Spaces. Cardiovasc Intervent Radiol 28, 561–569 (2005).

13. Wang, C. Y., Hu, J., Sheth, R. A. & Oklu, R. Emerging Embolic Agents in Endovascular Embolization: An Overview. Prog Biomed Eng (Bristol) 2, 012003 (2020).

14. Jensen, M. M. et al. Protein-based polymer liquid embolics for cerebral aneurysms. Acta Biomaterialia 151, 174–182 (2022).

15. Lord, J., Britton, H., Spain, S. G. & Lewis, A. L. Advancements in the development on new liquid embolic agents for use in therapeutic embolisation. J. Mater. Chem. B 8, 8207–8218 (2020).

16. Karadeli, H. H. & Kuram, E. Single Component Polymers, Polymer Blends, and Polymer Composites for Interventional Endovascular Embolization of Intracranial Aneurysms. Macromol Biosci 24, e2300432 (2024).

17. Rana, M. M. & Melancon, M. P. Emerging Polymer Materials in Trackable Endovascular Embolization and Cell Delivery: From Hype to Hope. Biomimetics (Basel) 7, 77 (2022).

18. Rodriguez, J. N. et al. Design and biocompatibility of endovascular aneurysm filling devices. J Biomed Mater Res A 103, 1577–1594 (2015).

19. Lord, J., Britton, H., Spain, S. G. & Lewis, A. L. Advancements in the development on new liquid embolic agents for use in therapeutic embolisation. J. Mater. Chem. B 8, 8207–8218 (2020).

20. Zalipsky, S. & Harris, J. M. Introduction to Chemistry and Biological Applications of Poly(ethylene glycol). in Poly(ethylene glycol) vol. 680 1–13 (American Chemical Society, 1997).

21. Knop, K., Hoogenboom, R., Fischer, D. & Schubert, U. S. Poly(ethylene glycol) in Drug Delivery: Pros and Cons as Well as Potential Alternatives. Angewandte Chemie International Edition 49, 6288–6308 (2010).

22. Veronese, F. M. & Pasut, G. PEGylation, successful approach to drug delivery. Drug Discov Today 10, 1451–1458 (2005).

23. Harris, J. M. & Chess, R. B. Effect of pegylation on pharmaceuticals. Nat Rev Drug Discov 2, 214–221 (2003).

24. Roberts, M. J., Bentley, M. D. & Harris, J. M. Chemistry for peptide and protein PEGylation. Advanced Drug Delivery Reviews 64, 116–127 (2012).

25. Harris, J. M., Martin, N. E. & Modi, M. Pegylation. Clin Pharmacokinet 40, 539–551 (2001).

26. Lin, C.-C. & Anseth, K. S. PEG Hydrogels for the Controlled Release of Biomolecules in Regenerative Medicine. Pharm Res 26, 631–643 (2009).

27. Cao, H., Duan, L., Zhang, Y., Cao, J. & Zhang, K. Current hydrogel advances in physicochemical and biological response-driven biomedical application diversity. Sig Transduct Target Ther 6, 426 (2021).

28. Zalipsky, S. Chemistry of polyethylene glycol conjugates with biologically active molecules. Advanced Drug Delivery Reviews 16, 157–182 (1995).

29. Arpicco, S. et al. Novel Poly(ethylene glycol) Derivatives for Preparation of Ribosome-Inactivating Protein Conjugates. Bioconjugate Chem. 13, 757–765 (2002).

30. Roberts, M. J., Bentley, M. D. & Harris, J. M. Chemistry for peptide and protein PEGylation. Adv Drug Deliv Rev 54, 459–476 (2002).

31. Greenwald, R. B., Choe, Y. H., McGuire, J. & Conover, C. D. Effective drug delivery by PEGylated drug conjugates. Advanced Drug Delivery Reviews 55, 217–250 (2003).

32. Akinc, A., Thomas, M., Klibanov, A. M. & Langer, R. Exploring polyethylenimine-mediated DNA transfection and the proton sponge hypothesis. The Journal of Gene Medicine 7, 657–663 (2005).

33. Chen, Z., Lv, Z., Sun, Y., Chi, Z. & Qing, G. Recent advancements in polyethyleneimine-based materials and their biomedical, biotechnology, and biomaterial applications. J. Mater. Chem. B 8, 2951–2973 (2020).

34. Chiu, D. T. et al. Small but Perfectly Formed? Successes, Challenges, and Opportunities for Microfluidics in the Chemical and Biological Sciences. Chem 2, 201–223 (2017).

35. Kilbride, B. F. et al. MRI-guided endovascular intervention: current methods and future potential. Expert Review of Medical Devices 19, 763–778 (2022).

36. Ma, Y. et al. Real-time x-ray fluoroscopy-based catheter detection and tracking for cardiac electrophysiology interventions. Medical Physics 40, 071902 (2013).

37. Zelepukin, I. V. et al. Flash drug release from nanoparticles accumulated in the targeted blood vessels facilitates the tumour treatment. Nat Commun 13, 6910 (2022).

38. Davis, F. J. & Mitchell, G. R. Polyurethane Based Materials with Applications in Medical Devices. in Bio-Materials and Prototyping Applications in Medicine (eds Bártolo, P. & Bidanda, B.) 27–48 (Springer US, Boston, MA, 2008). doi:10.1007/978-0-387-47683-4_3.

39. Blanco, P. J., Müller, L. O. & Spence, J. D. Blood pressure gradients in cerebral arteries: a clue to pathogenesis of cerebral small vessel disease. Stroke Vasc Neurol 2, (2017).

40. Jiménez, L. Á. C. et al. Model of Arteriovenous Malformation Created in Human Placenta for Training in Vascular Microneurosurgery. Oper Neurosurg 28, 418–426 (2025).

41. Burel, J. et al. The human placenta as a model for training and research in mechanical thrombectomy: Clarifications and use of the chorionic plate veins. Frontiers in Neurology 13, (2022).

42. Shimada, K., Kato, H., Saito, T., Matsuyama, S. & Kinugasa, S. Precise measurement of the self-diffusion coefficient for poly(ethylene glycol) in aqueous solution using uniform oligomers. J. Chem. Phys. 122, 244914 (2005).

43. Bird, R. B., Stewart, W. E., & Lightfoot, E. N. (1960). Transport Phenomena. John Wiley & Sons.

44. Modak, P., Hammond, W., Jaffe, M., Nadig, M. & Russo, R. Dynamic, 3D Schiff base networks for medical applications. Journal of Applied Polymer Science 137, 49756 (2020).

