## Supporting Information for "Soft-Robotic Magnetic Microfluidic Catheter for Delivery of Aqueous-Based Dual-Component Embolic Formulations"

### Microfluidic soft-robotic catheter for in-situ delivery of a personalized two-component hydrogel

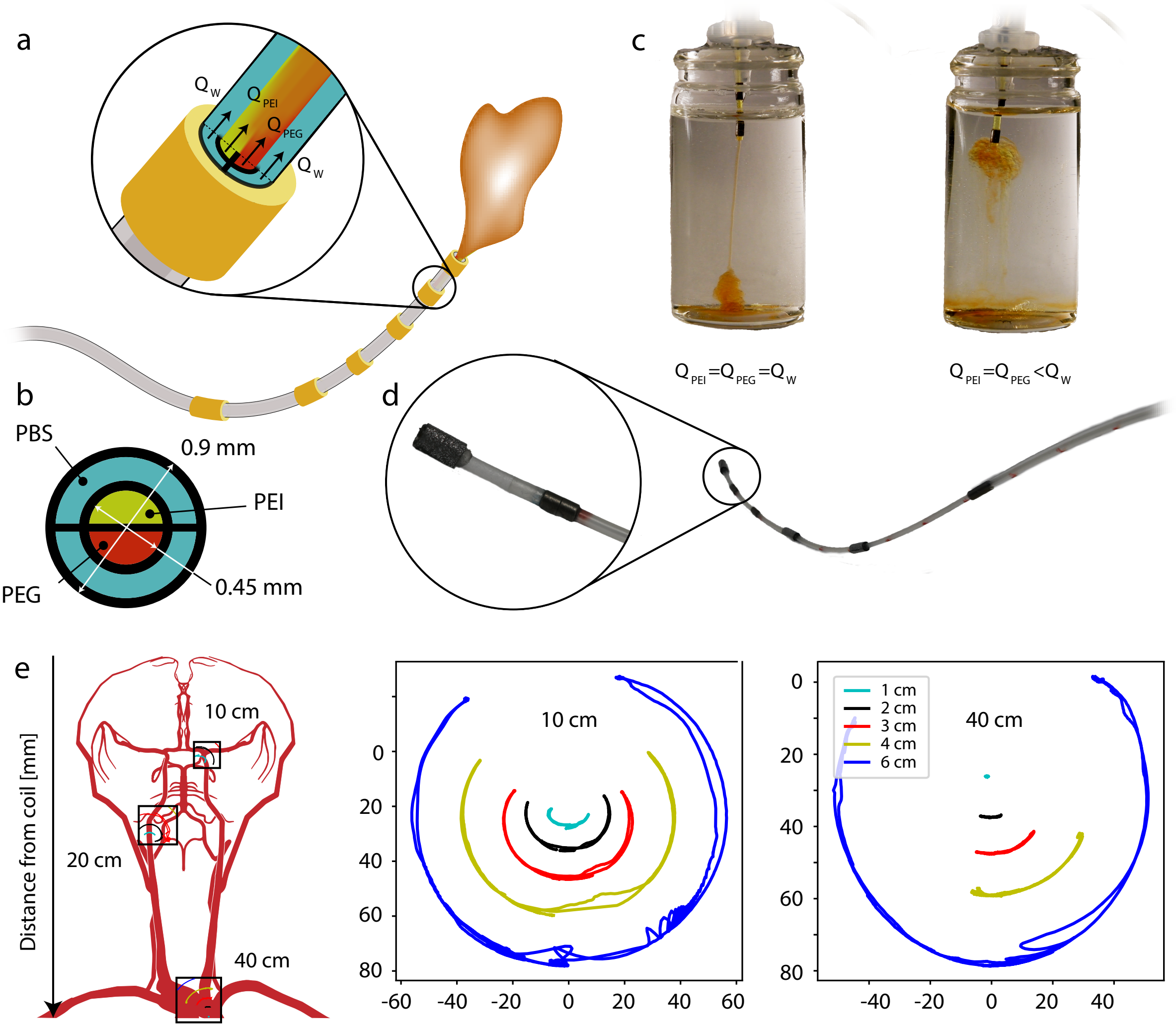


**Supplementary Fig. 1. Magnetic microfluidic catheter design and characterization.** (a) The overall design of the magnetic catheter with a series of magnets on the catheter surface and a microfluidic mixing chamber positioned at the tip of the catheter. (b) Cross section of the microfluidic catheter with a channel for PEI and PEG surrounded by two channels used for the injection of PBS and/or contrast agents. (c) Morphological properties of the injected embolic material based on the relative flow rates of the individual components. (d) Image of a microfluidic magnetic delivery catheter prototype with a zoomed in vision on the mixing chamber at the tip of the device. (e) Steerability and delectability of the magnetic catheter at various distances from the magnetic field generator surface.

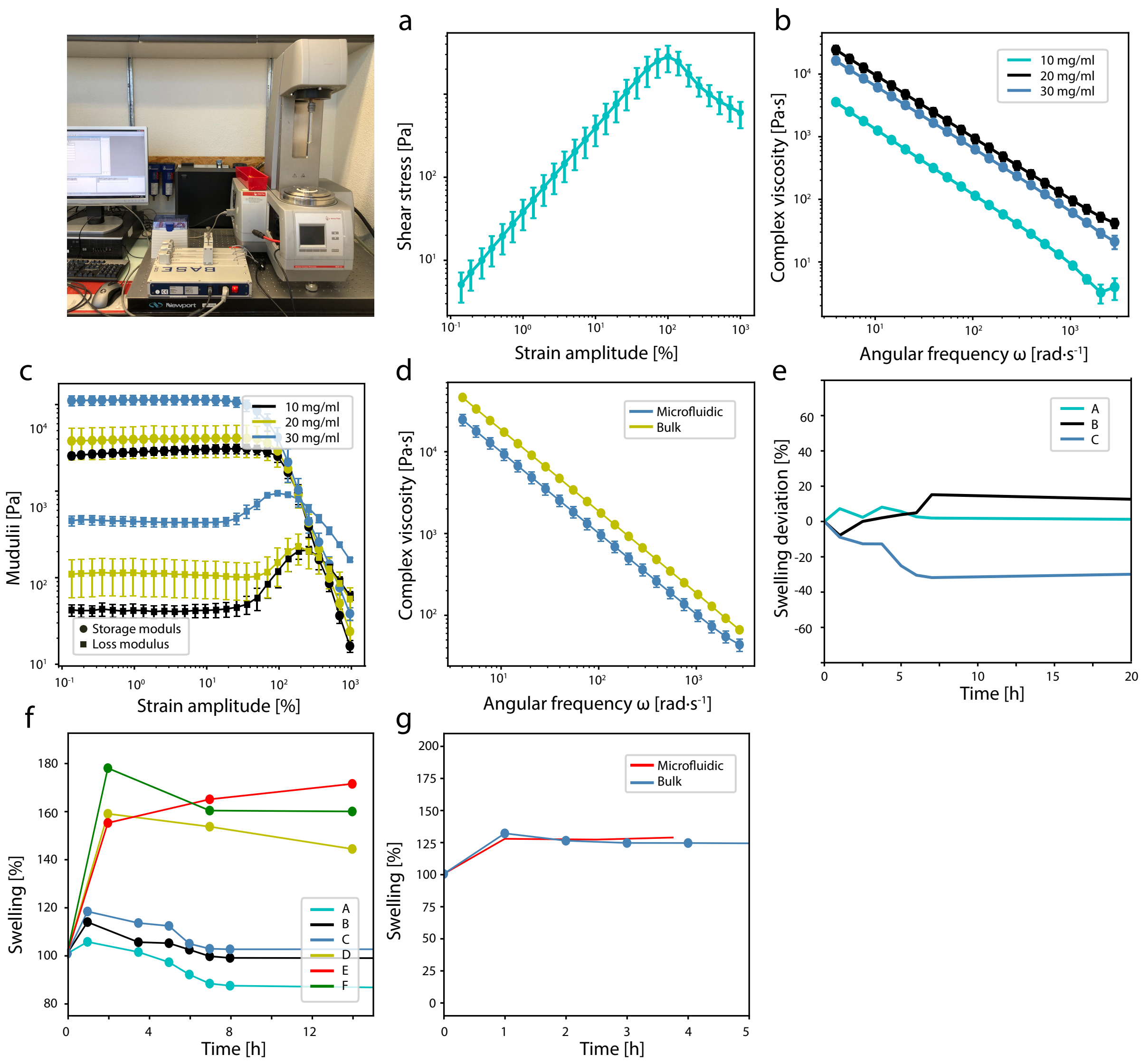


**Supplementary Fig. 2. Mechanical characterization of PEI-PEG hydrogels.** (a) Strain amplitude sweep. (b) Frequency sweep. (c) The storage and loss modulus of PEI-PEG hydrogels with varying concentrations of PEI. (d) Complex viscosity with various delivery methods. (e) Swelling properties of PEI-PEG hydrogels at different PEG concentrations with 40w% of Histodenz contrast medium relative to the unloaded hydrogel. (f) Swelling properties of PEI-PEG hydrogels at different PEI and PEG concentrations. (g) Swelling properties of PEI-PEG hydrogels with various delivery methods.

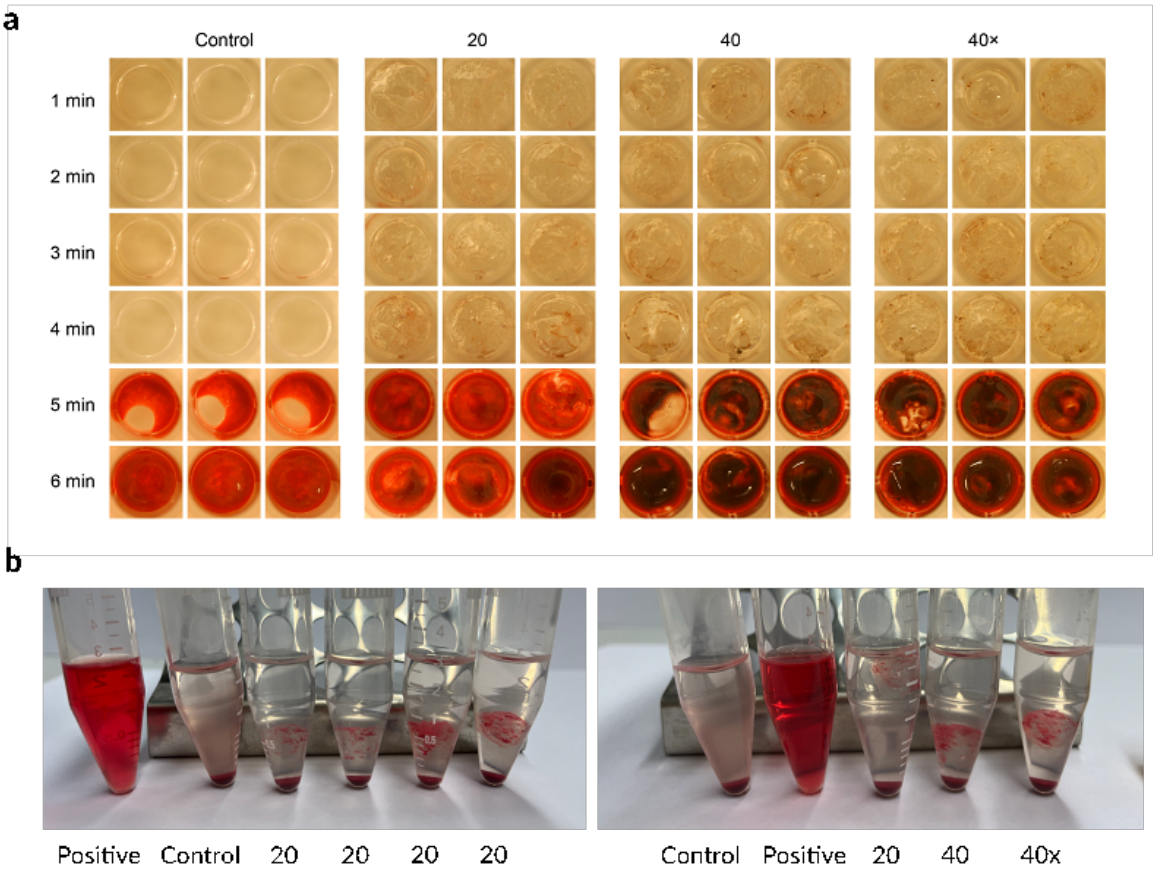


**Supplementary Fig. 3.** Biological characterization of PEI-PEG hydrogels containing. (a) The clotting time is unaffected when incubated with porcine blood. Blood without additives is used as a negative control. (b) Hemolysis results for different PEI concentrations and Histodenz loading suggest hemocompatibility of all materials.

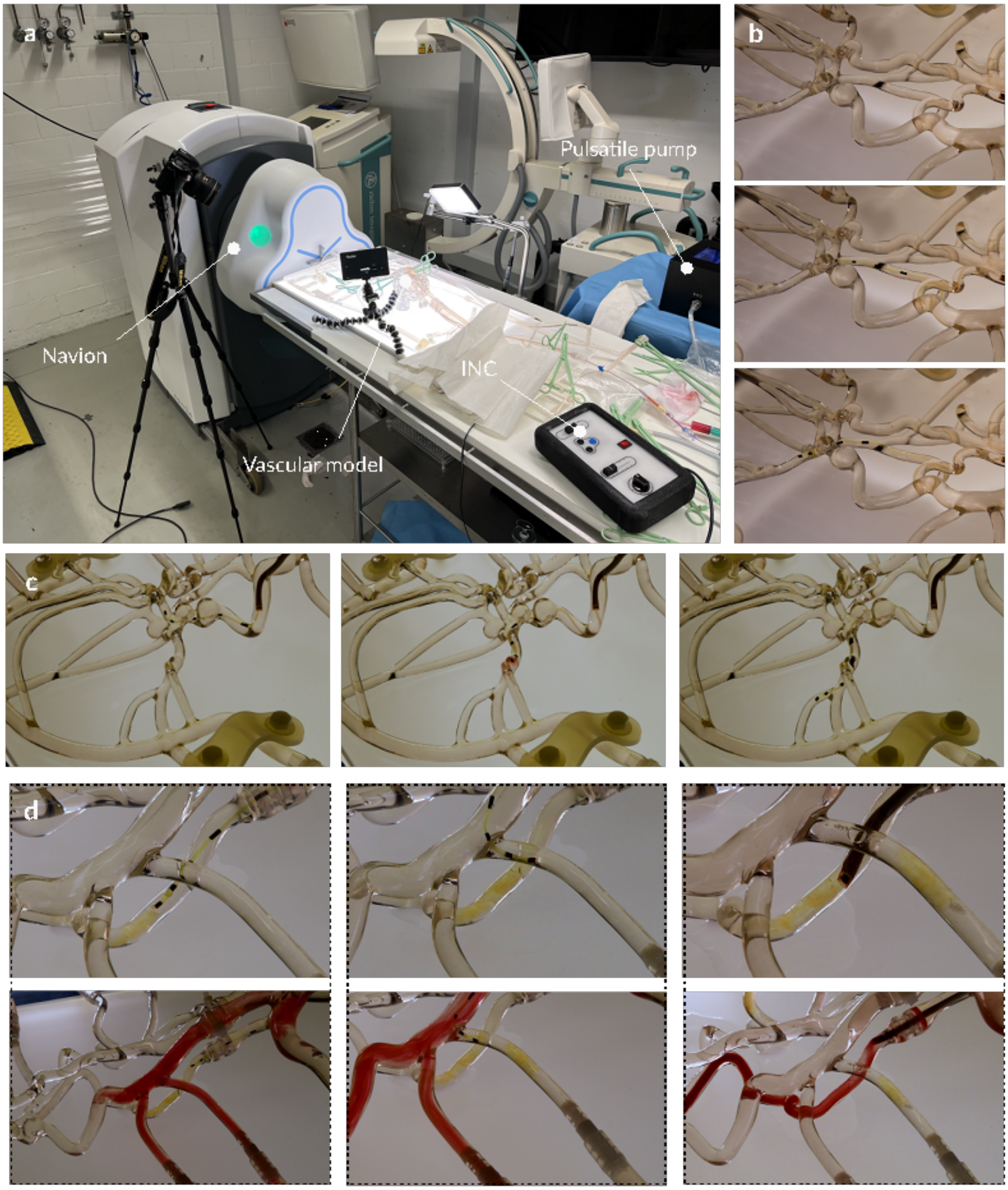


**Supplementary Fig. 4. Transcatheter embolization in in vitro human vasculature model.** (a) Experimental setup with silicone vascular model perfused with a pulsatile flow mimicking physiological conditions. (b) Navigation of the magnetic microfluidic catheter into the brain vasculature with a support catheter placed in the right vertebral artery. (c) Navigation of the magnetic microfluidic catheter into the brain vasculature with a support catheter placed in the right internal carotid artery. (d) TCE in the perfused right vertebral artery (left), right external carotid artery (middle), and aspiration of the embolic material in the right vertebral artery (right).

The lower images are showing the perfusion of the model after each embolic material injection or aspiration.

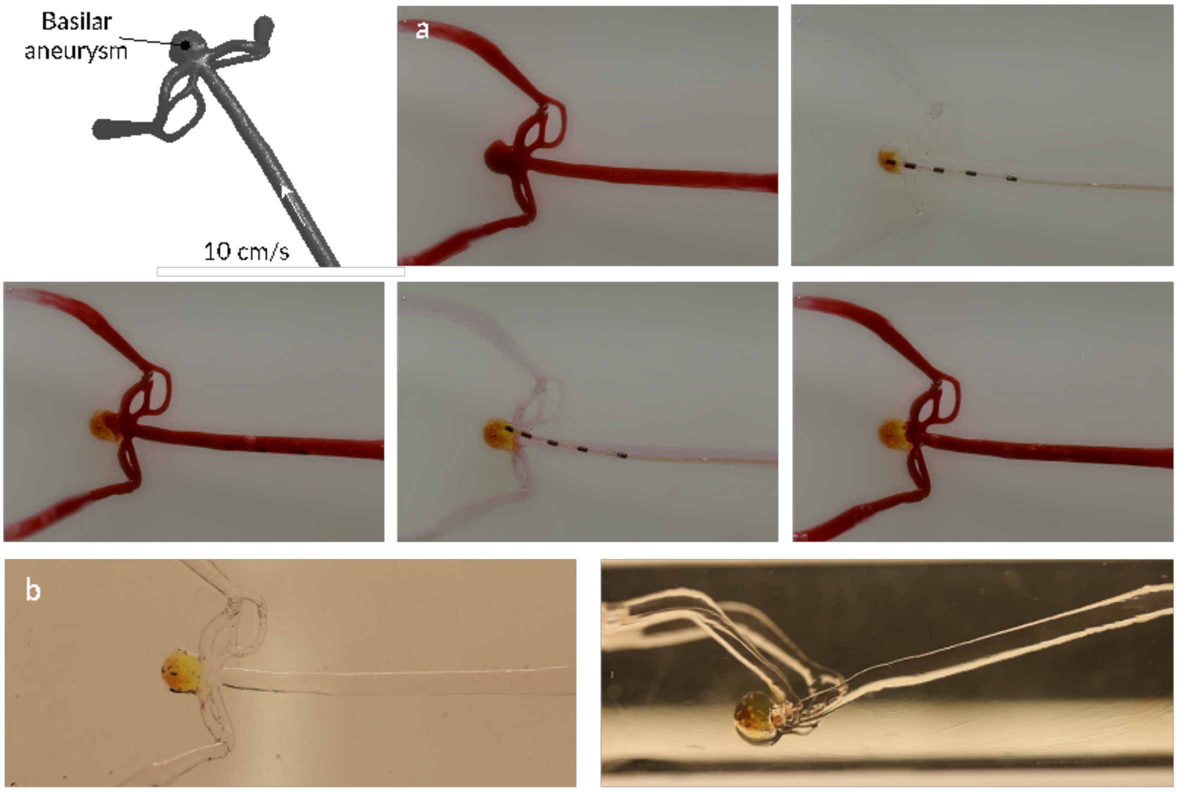


**Supplementary Fig. 5. Transcatheter embolization in in vitro basilar aneurysm models.** (top left) Schematic illustration of aneurysm model, including the perfusion parameters. (a) TCE sequence with full perfusion in the beginning followed by a first injection and partial perfusion of the aneurysm. Finally, it is followed by a complete filling without perfusion immediately after the injection. (b) Frontal and lateral images of the embolized model showing successful filling without residues outside of the aneurysm sac.

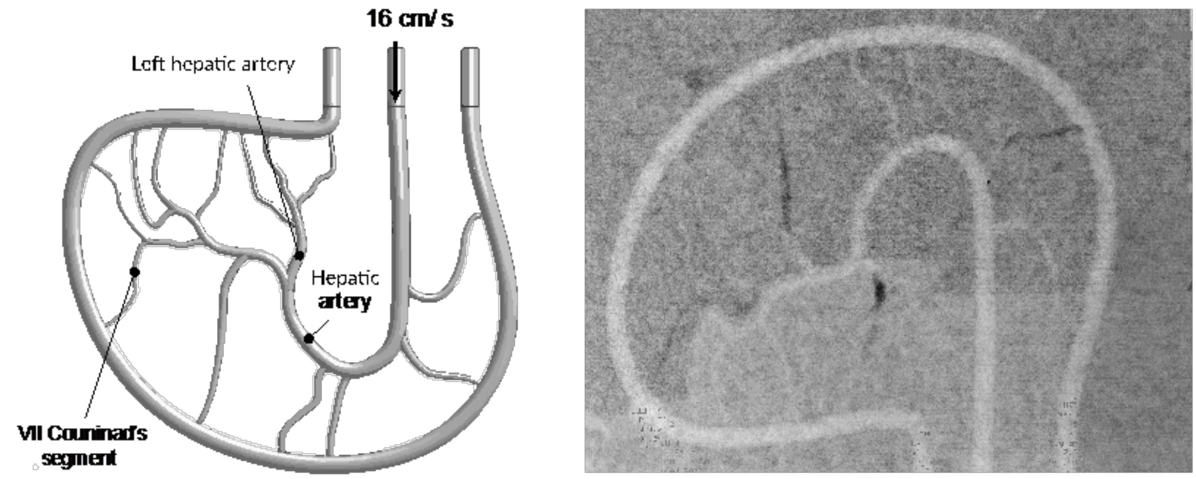


**Supplementary Fig. 6. Transcatheter embolization in in vitro liver model.** (left) Schematic illustration of the liver model, including the perfusion parameters. (right) Post-embolization fluoroscopic imaging shows good visibility of the embolization material both in small and large vessels.

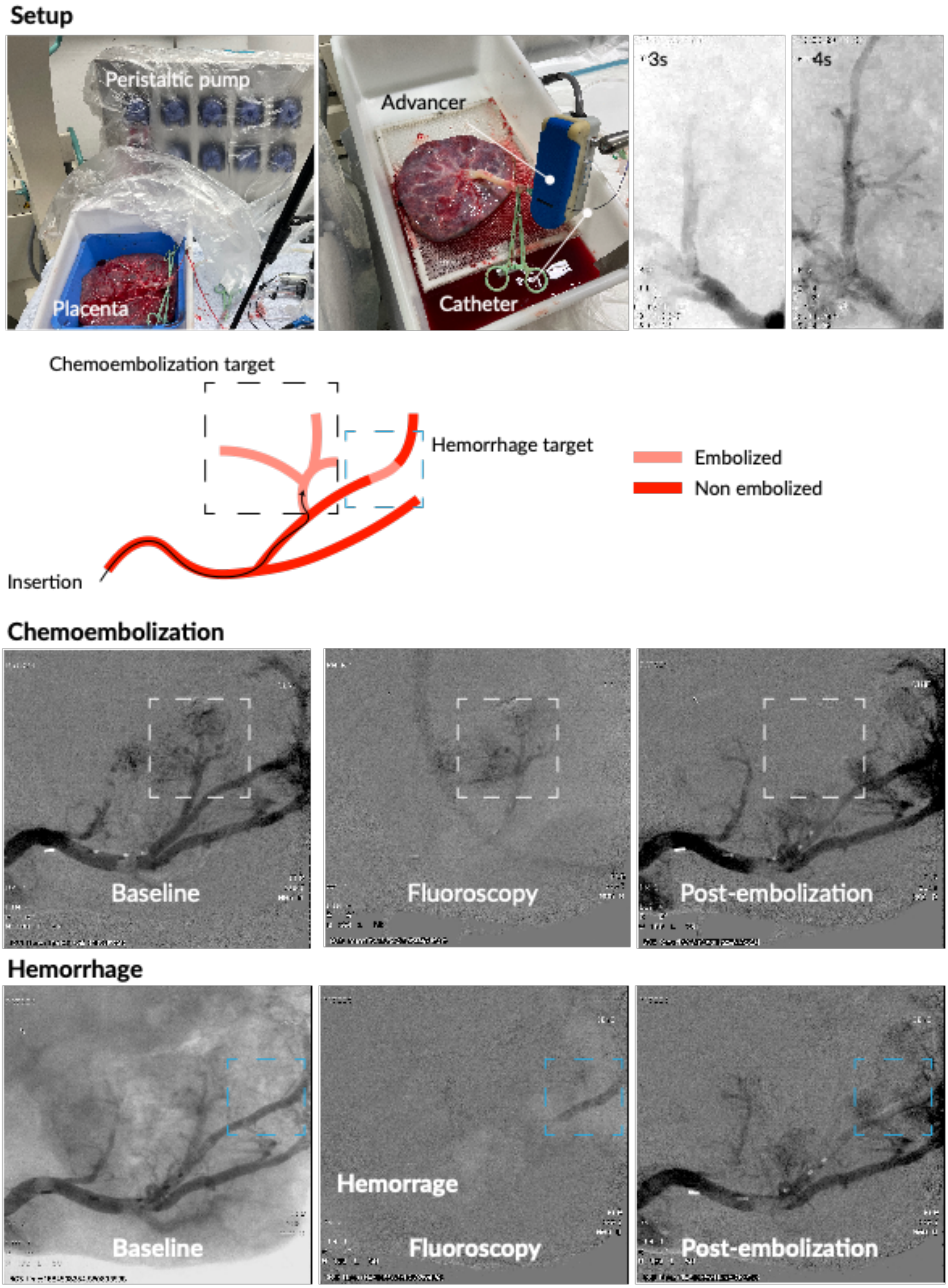
Setup
Peristaltic pump
Placenta
Advancer
Catheter
3s
4s
Chemoembolization target
Hemorrhage target
Embolized
Non embolized
Insertion
Chemoembolization
Baseline
Fluoroscopy
Post-embolization
Hemorrhage
Baseline
Hemorrage
Fluoroscopy
Post-embolization

**Supplementary Fig. 7. Selective embolization of sub-branches in an ex-vivo human placenta.** (Setup) An experimental setup illustrates the human placenta perfused by a peristaltic pump that mimics physiological flow conditions. The placenta is catheterized in the umbilical vein using an Avanti introducer sheet. The magnetic catheter is controlled with a mechanical advancer connected to the introducer sheet. Subbranches of the human placenta were selectively embolized for chemoembolization and hemorrhage treatment applications. Fluoroscopic image sequences show pre-embolization perfusion, injected embolic material, and post-embolization perfusion.

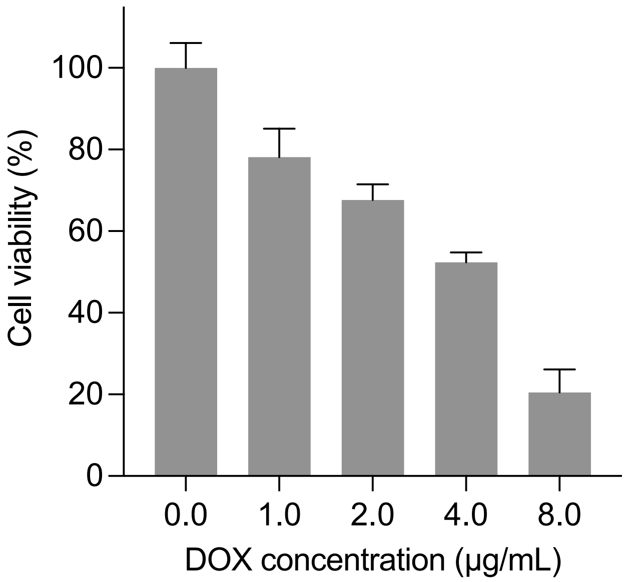


**Supplementary Fig. 8. Inhibitory Effects on Luciferase-Labelled Rat Liver Cancer Cells (CBRH-7919-luc) Post Incubation with PEI-PEG Hydrogel.** This figure illustrates the cell viability assessed by the MTT assay after a 48-hour treatment period with the PEI-PEG hydrogel (n = 3, data are presented as mean ± s.e.m.).

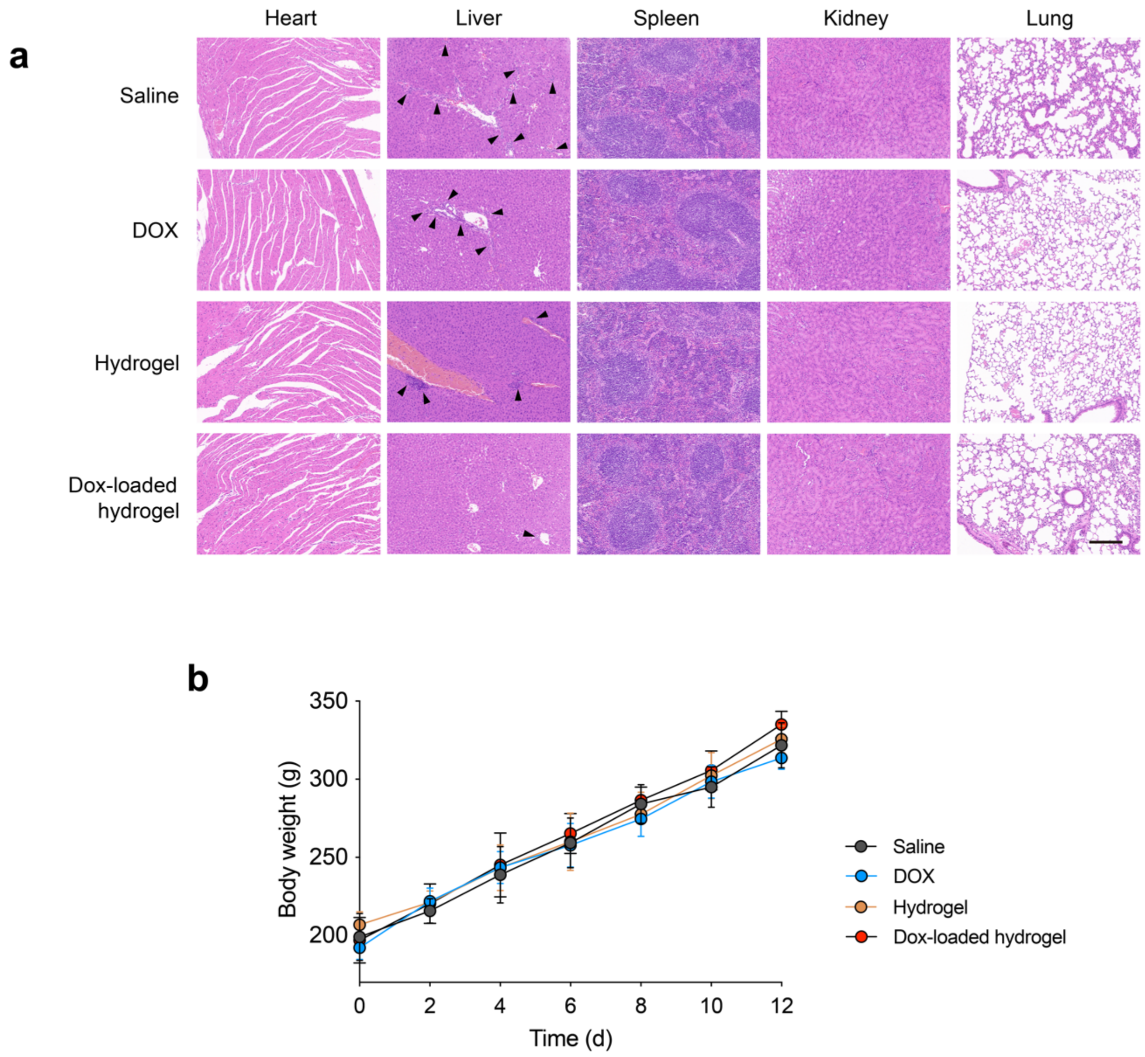


**Supplementary Fig. 9.** (a) Hematoxylin and eosin (H&E) staining of representative organ sections at day 12 post-delivery, focusing on major organs. Black arrows indicate regions of liver tumor infiltration. Scale bar represents 1 mm. (b) Graph depicting the changes in body weight of rats over the study period (n = 5), demonstrating the impact of the hydrogel treatment on systemic health.

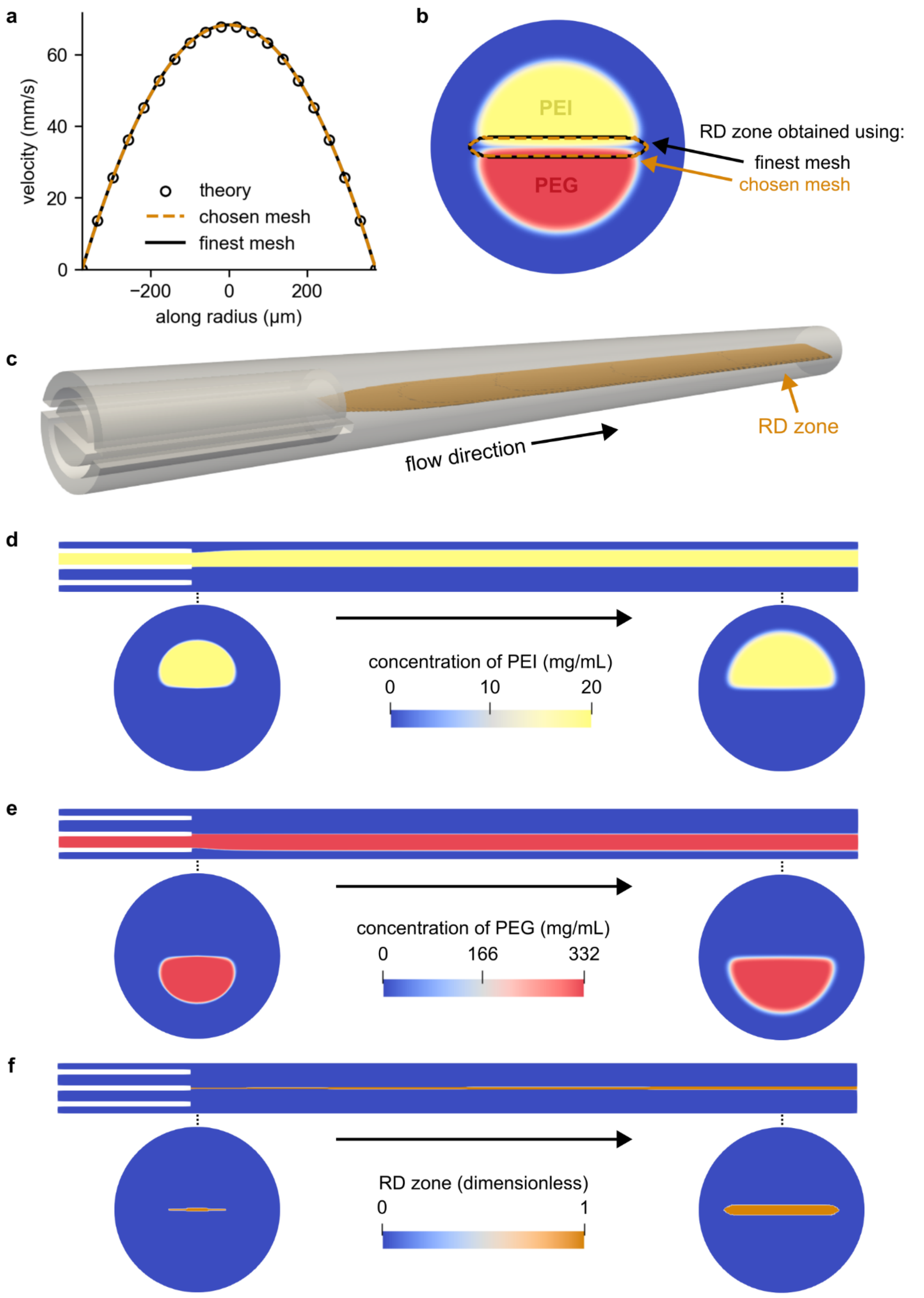


**Supplementary Fig. 10.** (a) Validation of numerical simulations and mesh testing, by plotting the theoretical velocity profile together with that obtained using different meshes. (b) Reaction-diffusion (RD) zone obtained at the outlet of the device from numerical simulations performed using different meshes. (c) Representation of the geometry of the multi-lumen microfluidic catheter device used in numerical simulations, and the RD zone (orange) forming along the device. (d) Concentration of PEI along the device. (e) Concentration of PEG along the device. (f) RD zone along the device.

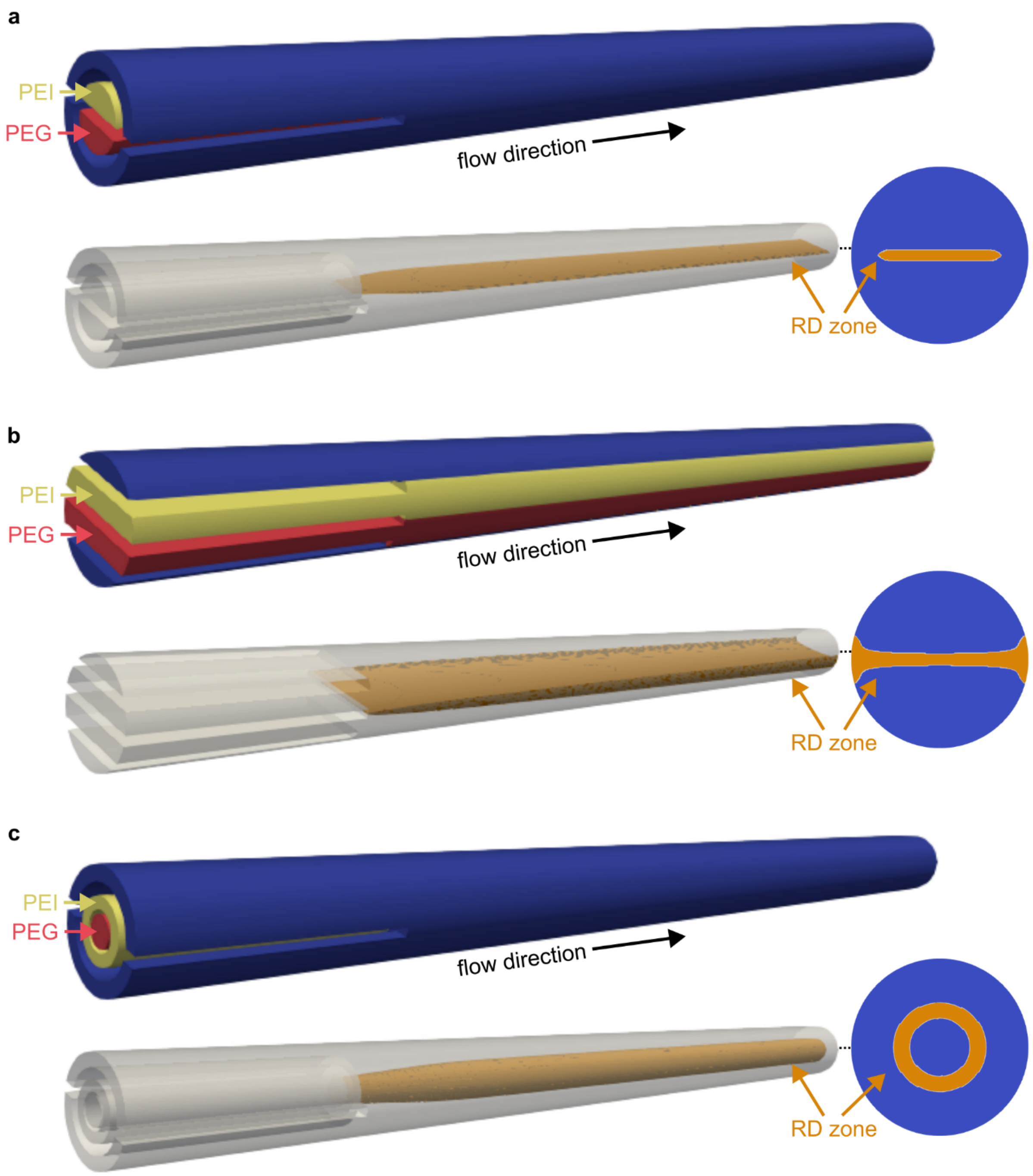


**Supplementary Fig. 11. Numerical simulation of various microfluidic catheter devices, performed to choose the optimal setup for PEI-PEG hydrogel formation.** (a) Multi-lumen microfluidic device. (b) 2D planar microfluidic device. (c) Co-axial microfluidic device. In each panel, we represent the location where PEI and PEG solutions are introduced, and the RD zone along the device and at the outlet.

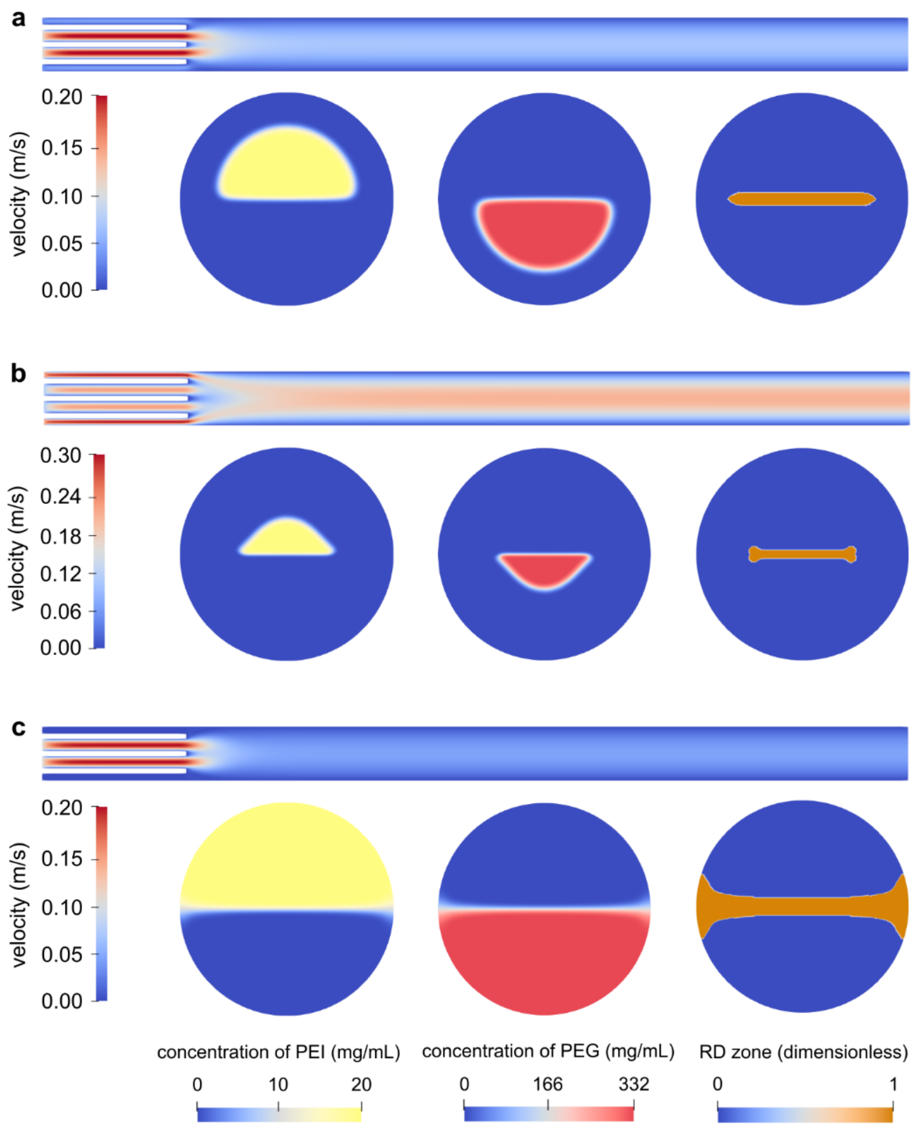


**Supplementary Fig. 12. Numerical simulation of the flow and mass transport in the multi-lumen microfluidic device considering various flow conditions**. (a) 300 µL/min for all flow rates (total flow rate = 900 µL/min), (b) sheath flow rate of 2000 µL/min (total flow rate = 2600 µL/min), (c) no sheath flow (total flow rate = 600 µL/min).

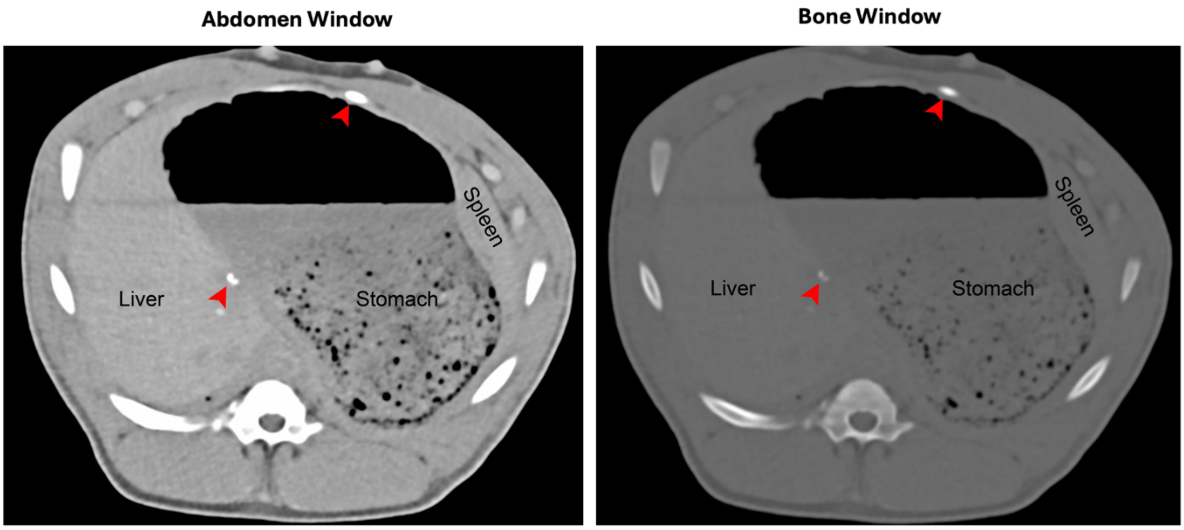


**Supplementary Fig. 13. Axial CT images in the venous phase following embolization of the splenic and hepatic veins.** The image on the left is displayed in the abdomen window, and the image on the right in the bone window. Red arrowheads indicate the embolization materials localized within the splenic and hepatic veins.

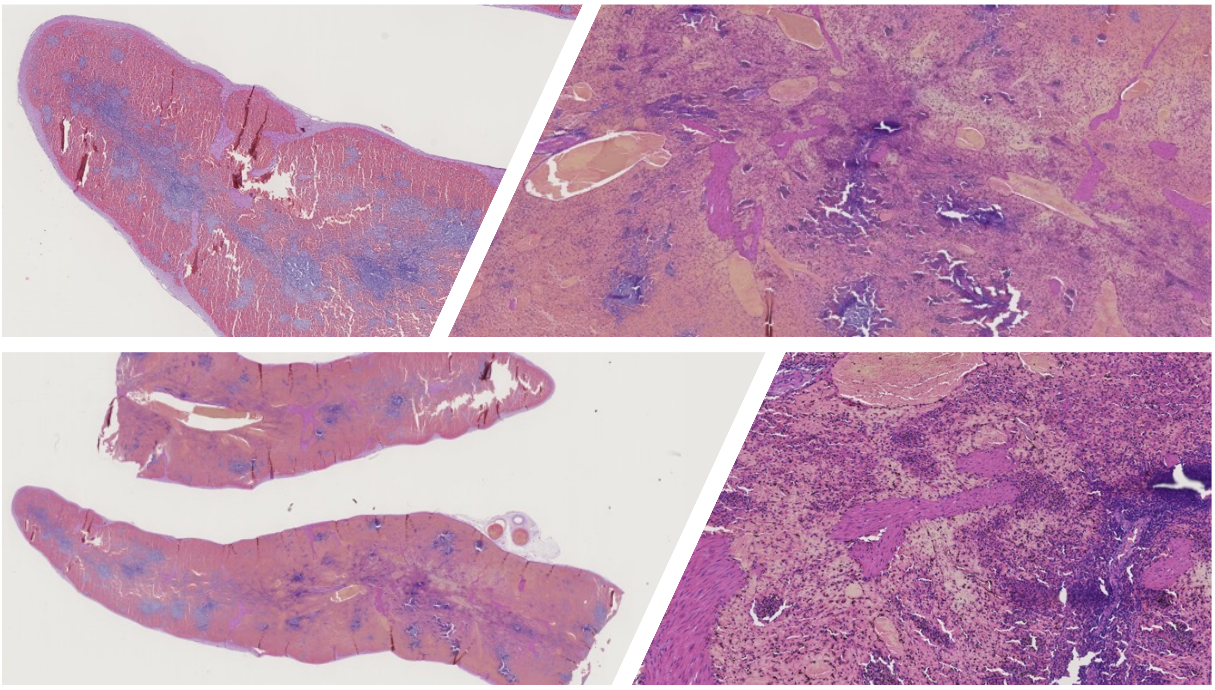

**Supplementary Fig. 14. Histopathological assessment of porcine spleen after embolization.** Representative H&E-stained sections of the spleen collected post-procedure. Low-magnification views (left) show preserved splenic architecture, while higher-magnification images (right) confirm the absence of abnormal tissue alterations, inflammatory infiltrates, or procedure-related damage in the examined regions.

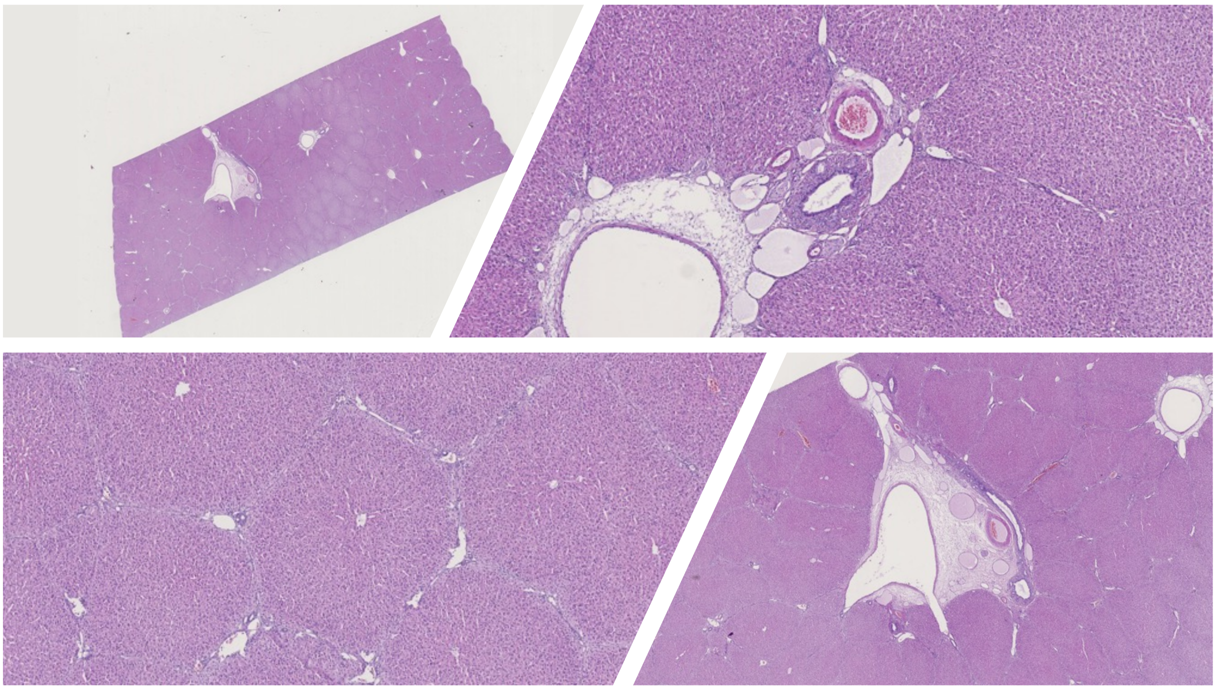

**Supplementary Fig. 15. Histopathology of porcine liver following embolization.** Representative H&E-stained liver sections obtained post-procedure at low and higher magnification, showing preserved tissue architecture and normal-appearing portal structures in the analyzed areas, without signs of overt inflammatory reaction or procedure-related damage.
